# Host and Microbe Scale Processes Shape Spatial Variation in *Aphaenogaster* (Hymenoptera: Formicidae) Genetics and Their Microbiota

**DOI:** 10.1101/2025.08.11.669684

**Authors:** Daniel A. Malagon, Benjamin Camper, Sophie Millard, Curt Harden, Ernesto Recuerdo Gil, Michael Caterino, Maslyn Ann Greene, Anna Seekatz, Seth R. Bordenstein, Sarah R. Bordenstein, Sharon Bewick

## Abstract

Like all ecological communities, host-associated (HA) microbiota are shaped by environmental selection and dispersal limitation. However, unlike communities of free-living organisms, communities of HA microbes experience selection and dispersal at two separate scales – the scale of the microbes and the scale of their hosts. Thus, HA microbes must tolerate not only the environment created by their host (microbe-scale environment), but also, the environment in which their host resides (host-scale environment). Likewise, HA microbes can disperse between hosts through either horizontal or vertical transmission (microbe-scale dispersal) but can also disperse between locations through host movement (host-scale dispersal). In this paper, we examine how multiscale environmental selection and dispersal limitation shape the genetics and HA microbiota of ants in the *Aphaenogaster fulva-rudis-texana* (Hymenoptera: Formicidae) complex. We begin by showing how spatial variation in *Aphaenogaster* genetics is shaped by host-scale environmental selection and dispersal limitation. We then show how this allows both host- and microbe-scale environmental selection to govern spatial variation in *Aphaenogaster* microbiota. Finally, we discuss the possibility that microbe-scale dispersal limitation also impacts spatial variation in *Aphaenogaster* microbiota and that this, in turn, may contribute to spatial variation in *Aphaenogaster* genetics. Ultimately, our results help to shed light on the myriad of interacting factors governing spatial variation in HA microbiota, including the potential for complex, bidirectional interactions between host- and microbe-scale processes.

## Introduction

Ecological communities vary across space.^1–3^ Two of the most important drivers of spatial variation are environmental selection and dispersal limitation.^4^ Environmental selection (i.e., genotype and/or species sorting) causes spatial structure by limiting access of certain taxa to certain locations based on their ability to tolerate local biotic or abiotic conditions. This can yield a wide array of different spatial patterns, including the evolution of ecotypes, genetic isolation by environment (IBE),^5,6^ ecotones, ecoclines and distance-decay of community similarity.^7^ Like environmental selection, dispersal limitation also causes spatial structure by limiting access of certain taxa to certain locations.^8^ In the case of dispersal limitation, however, access is limited by the ability of taxa to cross physical barriers and/or unsuitable habitat.^8^ Similar to environmental selection, dispersal limitation can result in a wide variety of different spatial patterns, including range boundaries, genetic isolation by distance (IBD), metapopulations, metacommunities and, once again, distance-decay of community similarity.

Like all ecological communities, host-associated (HA) microbiota exhibit spatial variation in taxon community composition.^9,10^ However, whereas spatial structure in communities of free-living organisms is primarily governed by environmental selection and dispersal limitation at the scale of the organism, spatial structure in HA microbes is governed by environmental selection and dispersal limitation at two scales: the scale of the microbe and the scale of their host. Thus, spatial variation in HA microbes is affected not only by dispersal of microbes from one host to another (microbe scale dispersal limitation) and differences in conditions on the host (microbe scale environmental selection; e.g., due to differences in host genetics), but also by dispersal of hosts from one location to another (host scale dispersal limitation) and differences in conditions where the hosts reside (host scale environmental selection).^9,11^

Just as host scale processes can impact the spatial ecology of HA microbiota (i.e., top-down effects), microbe scale processes can influence the spatial ecology of their hosts (i.e., bottom-up effects).^13^ Many microbes, for instance, alter host dispersal. In some cases, this is an indirect effect of microbes negatively impacting host health and, as a result, reducing dispersal ability (e.g., hosts displaying signs of a microbial illness often disperse less or not at all).^14^ In other cases, however, microbes have actually evolved to manipulate host dispersal, often in ways that promote their own transmission. Classic examples include *Cordyceps* spp. that cause ants to move towards the tops of trees^15^ where they can be eaten by birds and *Toxoplasma gondii* that causes mice to be attracted to cat urine.^16^ In addition to host dispersal, HA microbes can also alter host environmental selection. In some systems, for example, it is the abiotic tolerances of obligate endosymbionts, rather than the hosts themselves that restrict environmental associations.^17^ In other systems, hosts are limited by their own environmental constraints, but microbial symbionts can positively affect these constraints by increasing host tolerance^17^ or altering host resource use.^18^ This then allows hosts to occupy niches that they cannot in absence of the symbiont.

Beyond ecology, HA microbiota can also impact host spatial patterns by affecting host evolution. A variety of HA microbial taxa, for instance, manipulate host reproduction, impacting patterns of gene flow within and between lineages.^19^ This can alter effective migration rates between populations, causing the emergence of spatial structure in the host where it would not otherwise occur.^20^ Examples of reproductive manipulators include microbes that induce cytoplasmic incompatibility (CI), male-killing, feminization, and parthenogenesis to increase their fitness.^21^ Perhaps the best-known and most widespread example is the genus *Wolbachia*. Present in almost half of all arthropod microbiota,^22^ this bacterium imposes a myriad of effects on its host,^21,23^ including induction of CI among hosts harboring differing *Wolbachia* strains or even different loads of the same strains.^24,25^ CI can then lead to post-zygotic reproductive barriers that prevent gene flow between hosts, often in a spatially explicit manner.^26,27^

The bilateral dynamics between a host and its microbiota ultimately shape organization of the host and microbiota consortium. This has led to the development of a lexicon and disciplinary matrix for holobiont biology.^12^ However, while the various top-down and bottom-up processes that govern relationships between HA microbiota and their hosts are well-known, what is less well understood is how these processes combine to determine spatial variation. This includes spatial variation in hosts, in HA microbes, in entire HA microbiota and in the holobiont as a whole. In this study, we use *Aphaenogaster fulva-rudis-texana* (Hymenoptera: Formicidae; henceforth ‘*A. rudis*’) ant complex ants in Great Smoky Mountains National Park (GSMNP) to investigate the role of host and microbe scale spatial processes on spatial patterns in *Aphaenogaster* population genetics and *Aphaenogaster-*associated microbiota. Using previously derived, high resolution GSMNP bioclimatic variables,^28,29^ we assess the extent to which host genetics and HA microbiota vary as a function of the environment of the host (i.e., host scale environmental selection) and spatial distance (i.e., host scale dispersal limitation). We then assess the extent to which HA microbiota vary as a function of host genetics (i.e., microbe scale environmental selection). Finally, we examine the role of microbe scale dispersal limitation on endosymbionts – a subset of HA microbial taxa that are more likely to exhibit microbe scale dispersal limitation than other members of the *Aphaenogaster* microbiota. We hypothesize that both host and microbe scale environmental selection will impact spatial variation in HA microbiota. By contrast, we hypothesize that the high mobility of *Aphaenogaster* alates will minimize the role of host scale dispersal limitation.^30^ We do, however, expect to see evidence of microbe scale dispersal limitation among endosymbionts.

## Materials and Methods

### Study Species

Ants of the *A. rudis* complex are abundant and widespread in eastern hardwood forests of North America.^31^ With a generalist strategy that includes colonization of a wide array of forest types,^32^ and broad latitudinal,^33^ and elevational ranges,^34–38^ this complex is ideal for examining effects of host scale environmental selection on HA microbiota. Further, *A. rudis* complex ants within GSMNP comprise four incipient species - *A. rudis*, *A. picea*, *A. fulva*, and *A. carolinensis*. Generally speaking, *A. picea* is found at higher elevations while *A. rudis* is found at lower elevations.^39^ Despite these differences,^39,40^ members of the complex co-occur extensively, both within GSMNP and across the Southeast broadly, ^41^ possibly because they are in the early stages of lineage divergence.^41^ This makes the *A. rudis* complex ideal for examining the effects of microbe scale environmental selection (i.e., host genetics) on HA microbiota.

### Study Sites

All ants were collected from GSMNP. GSMNP is an ideal location for studying the effects of host scale environmental selection due to its mountainous terrain. In particular, the large elevational gradients characteristic of the GSMNP landscape result in significant bioclimatic variation over relatively small distances.^28,29^ This means that there is wide variation in environmental selection over distances that are small relative to the dispersal capabilities of ants, allowing us to minimize the role of host scale dispersal limitation. *A. rudis* complex ants were collected from 12 All Taxa Biodiversity Index (ATBI) plots^42^ as well as an additional plot at one of the lowest elevations in the park (13 total sites) from Fall 2019 - Summer 2021. These plots vary greatly in elevation (522-1,494 meters) and were selected to represent the dominant ecosystems present in GSMNP. At each plot, we collected between 4 and 15 1m x 1m quadrats of litter across 6 to 11 unique time-points, depending on site accessibility. All quadrats within any sampling period were selected to be at least 5m apart to avoid sampling the same colony within multiple quadrats. In addition to quadrat sampling, we spent ∼30 minutes searching for and aspirating ants during each visit. Because of low capture rates in the summer and winter, along with a lack of significant difference between the microbiota of ants captured in the spring versus fall (see SI III S3.15), we removed ants sampled in the summer/winter from our analysis and pooled all ants collected in the spring/fall for subsequent analyses. (For additional details regarding ant collection and identification, see SI I).

### COI and CAD Gene Extraction, Amplification, and Sequencing

Genomic DNA was extracted from 1 leg of each ant using the Thermo Scientific Genejet Genomic DNA purification kit following the manufacturer’s protocol. We amplified the COI gene following Harden et al. 2022^48^ using the primers LCO1490 and HCO2198 in 25-µL PCR reactions containing 2.5 µL of template DNA, 1.0 µL of each primer, 2.5 µL of dNTPs, 2.0 µL of MgCl2, 0.125 µL of Platinum Taq, 2.5 µL of buffer, and 13.375 µL of Invitrogen ultra-pure water (Catalog number: 10977023). Reactions were carried out using an Eppendorf MasterCycler with the following settings: initial denaturation stage of 180 seconds at 95 °C followed by 35 cycles of a denaturation stage at 94 °C for 30 seconds, an annealing stage at 45 °C for 30 seconds, an extension stage at 72 °C for 45 seconds, and ending with a final extension at 72 °C for 180 seconds. CAD was amplified using primers CD439F and CD851R.^51^ CAD reactions were carried out as above using an Eppendorf MasterCycler beginning with an initial denaturation at 95°C for 3 minutes followed by 40 cycles of a denaturation stage at 94°C for 30 seconds, an annealing stage at 50 °C for 30 seconds, an extension stage at 72 °C for 50 seconds, and a final extension at 72 °C for 5 minutes. Amplified genes were forward and reverse Sanger sequenced at Psomagen.

### COI and CAD Phylogenetic Tree Construction

Forward and reverse COI and CAD Sanger sequencing reads were merged in R (v.4.2.1)^49^ using the package sangeranalyseR (v.1.6.1).^50^ After quality control, 91 CAD sequences and 80 COI sequences remained from our original 102 samples. We then aligned merged reads, without a reference, using our own data as well as data from an *A. umphreyi* individual that we chose as our outgroup (GenBank Accessions: KP860492.1 and KP730141.1).^47^ COI and CAD genes were aligned separately using the MAFFT algorithm in Mesquite. We then employed maximum likelihood methods for phylogenetic tree construction through the web server version (v. 2.4.0) of IQ-Tree^51^ using default parameters. This was repeated for the COI and CAD single alignments separately to produce single gene trees. For the COI (mtDNA marker) and CAD (nuDNA marker) genes, the best-fit models of nucleotide substitution, selected based on BIC and using ModelFinder within IQ-Tree, were TPM2u+F+R2 and K2P+R2 respectively. Tree discordance between COI and CAD gene trees was calculated using the *cospeciation* function and visualized using the *cophylo* function both in the *phytools* package in R (version 4.2.1). Because of the high level of discordance (see below) between COI and CAD trees, along with the fact that *Wolbachia* (see below) often impacts mtDNA inheritance patterns differently from the inheritance of nuclear DNA, we chose to perform all statistics using separate COI and CAD trees rather than a concatenated tree. For host-specific analyses, we used all COI and/or CAD sequences available (80 and 91 samples respectively). For analyses that paired COI/CAD and microbiota, we only used samples for which both host and microbiota data were available (46 samples for COI/microbiota and 52 samples CAD/microbiota). For all analyses, we use a square-root transformation of branch lengths^52^ in either the COI or CAD trees.

### Microbiome Extraction, Amplification, and Sequencing

*Aphaenogaster* microbial DNA was extracted using the ZymoBIOMICS DNA Microprep kit (catalogue number D3401). Prior to extraction, individual ants were surface sterilized for 1 minute in 10% bleach solution and rinsed 3 times in ultra-pure water to ensure transient microbes on the exoskeleton were removed. We performed the extraction following the manufacturer’s protocol with the following modifications: 600 uL input of ant lysate was added to 1800 uL of binding buffer due to low input. We skipped filtering steps 4, 12, and 13 due to low input and heated the elution buffer to 60 °C prior to elution to maximize DNA concentrations. Samples were lysed for 40 minutes using a vortex genie and for 20 minutes using a tissuelyzer2 at 25 x/second to effectively lyse the ant exoskeleton.

To characterize bacterial communities we amplified the V4 region of the 16S rRNA gene using touchdown PCR with common dual index primers.^53^ We amplified in 20-µL PCR reactions containing 5 µL DNA, 5 µL of the primer set (515F and 806R), 2 µL of 10x Accuprime PCR Buffer II, 0.15 µL of Accuprime HiFi polymerase, and 7.85 µL of ultra-pure water per reaction. Reactions were carried out using an Eppendorf MasterCycler with the following settings: an initial denaturation stage of 120 seconds at 95 °C followed by 20 cycles of a denaturation stage at 95 °C for 20 seconds, an annealing stage at 60 °C for 20 seconds, and an extension stage at 72 °C for 5 minutes. This was followed by 20 cycles of a denaturation stage at 95 °C for 20 seconds, an annealing stage at 55 °C for 20 seconds, an extension stage at 72 °C for 5 minutes, and a final extension stage at 72 °C for 10 minutes. Samples were normalized prior to pooling using a SequalPrep Normalization Plate Kit (catalogue number #A1051001). Pooled sample DNA concentrations were quantified via KAPA qPCR. Multiplexed pooled libraries were sequenced paired end 2 x 300 cycles on an Illumina NextSeq 2000 to an average depth of 245,448 reads. Resulting FASTQ files were analyzed using a custom Qiime2^54^ pipeline. (For additional information on Taxonomic Assignment and Dataset Manipulation, see SI I; see also Github repository below).

### Endosymbionts

Thirty-two separate ASVs were taxonomically classified to the *Wolbachia* genus, including members of supergroups A, B and F (see SI I Figure S1.1).^21^ Only three of these ASVs, however, independently comprised more than 0.01% of reads across all ant microbiota (two from supergroup A, and one from supergroup F). These were also the only *Wolbachia* ASVs found in more than one ant. To compare *Wolbachia* relative abundance across host clades (see below), we pooled all *Wolbachia* ASVs to genus and compared relative abundances at the genus level. To perform ASRs and haplotype maps, however, we considered each of the three dominant *Wolbachia* ASVs independently, focusing on the presence/absence (P/A) of each strain on each ant. Specifically, we set the strain as ‘present’ if it comprised more than 0.5% of reads on an animal and ‘absent’ otherwise. We used a threshold of 0.5%, rather than 0%, because most samples had a small handful of *Wolbachia* reads from one or more of the three dominant *Wolbachia* strains. These reads could represent cross-contamination that was not removed during our cleaning steps (see above) or true presences at very low abundance. Regardless, there was a clear distinction between samples with only a few reads from dominant *Wolbachia* strains and samples with a large number of reads from dominant *Wolbachia* strains (see the ‘elbows’ in SI I Figure S1.2A-C). A threshold of 0.5% was selected because it separated the low versus high read samples for all three *Wolbachia* strains. Importantly, even if low read samples represent true presences, and not contamination, large differences in *Wolbachia* loads can lead to cytoplasmic incompatibility (CI) if a strain is too rare in females and their eggs to rescue sperm modification by high density males.^57^ Similar methods were used on three additional endosymbionts (*Spiroplasma*: 32 ASVs, 5 ASVs>0.01% of reads; *Entomoplasma*: 5 ASVs, 1 ASV>0.01% of reads; *Sulcia*: 1 ASV, 1ASV>0.01% of reads) to compare abundances across ants and to construct ASRs. However, because of the overall lower abundances of these endosymbionts, we only required that the endosymbiont be present at 0.1% of reads to classify as ‘present’ (see respective elbows in SI I Figure S1.2D-J).

### Statistical Analyses

All statistical analyses were performed in R (v.4.2.1). In the main text we focus on results using the COI tree and phylogenetically aware incidence metrics (Faith’s PD^58^ and Unifrac dissimilarity^59^). Abundance-based and/or phylogenetically unaware metrics as well as results with the CAD tree are reported in the SI. All data and code can be found in the following github repository https://github.com/dmalago/Apheanogaster_GSMNP_Microbiome.

#### Mantel tests

We used Mantel^60^ tests to assess whether host relatedness varied with geographic distance, whether host relatedness varied with environmental similarity, whether HA microbiota similarity varied with geographic distance, whether HA microbiota similarity varied with environmental similarity and whether HA microbiota similarity varied with host relatedness. All Mantel tests were performed using the *mantel* function from the *vegan* package(v. 2.6.4).^61^ We then used the *mantel.correlog* function, also from the *vegan* package (v. 2.6.4), to calculate mantel correlations at varying geographic or environmental distance classes or host relatedness classes. For spatial distances, we used the Haversine formula to calculate the great circle distance between capture locations based on longitude and latitude. This was implemented using the *distm* function from the *geosphere* package (v. 1.5-20). For environmental distance, we considered elevation, mean soil moisture (volumetric content), soil minimum temperature, and soil maximum temperature derived from previous GSMNP studies.^28,29^ We first z-transformed these four environmental variables, and then calculated environmental distances based on the Euclidean distances between the z-transformed environmental variable values at each capture location. For host relatedness, we used distances based on the branch lengths of the scaled COI or CAD trees. Branch lengths were calculated using the *cophenetic.phylo* function from the *ape* package (v. 5.7.1).^62^ For microbiota similarity we used Jaccard,^63^ Bray-Curtis,^64^ Unifrac,^59^ and weighted Unifrac indices.^59^ For several analyses, we also performed partial mantel tests, regressing microbiota similarity against both spatial and environmental distance. This was done using the *mantel.partial* function from the *vegan* package.

#### Ancestral state reconstructions (ASRs)

We performed ancestral state reconstructions (ASRs) for host environment using a one-dimensional ‘conglomerate’ environmental variable. Briefly, using the same four environmental variables we considered above, we performed a principal component analysis (PCA) based on the environmental variables associated with each ant. We then selected the first principal component (PC1) and used the value of each ant along this axis for our ASR. ASRs were generated using the *fastAnc* function from the *phytools* package (v. 2.3.0). We also performed ancestral state reconstruction for host carriage of each of the three dominant *Wolbachia* strains (see above), five dominant *Spiroplsama* strains, one dominant *Entomoplasma* strain and one dominant *Sulcia* strain. This was done using the *ace* function from the *ape* package.

#### Clade Comparisons

Using the COI consensus tree, we identified 2 main clades (henceforth the “blue” and “purple” clades) as well as a set of basal samples (henceforth the “red” (paraphyletic) clade). Using the same conglomerate environmental axis that was used for ASR (see above), we tested for differences in environmental conditions between clades using the *kruskal.test* function from the *stats* package (v. 4.2.1) followed by post-hoc testing with Mann-Whitney U tests using the *wilcox.test* function, also from the *stats* package. We similarly tested for differences in *Wolbachia* relative abundance (pooling all *Wolbachia* ASVs to genus) across clades, as well as differences in microbiota diversity, both including and excluding endosymbionts (see SI I).

#### Regression Analyses

We used generalized linear models^65^ with beta distributions to regress both *Wolbachia* relative abundances and diversity with and without endosymbionts against principal component one (PC1) of the environmental PCA (see above). This was done for all ants and then for the “blue”, “purple” and “red” clades individually. All regressions were performed using the *betareg* function from the betareg package.

#### Haplotype Networks

We constructed haplotype networks using the Templeton, Crandall, and Sing (TCS) network^66^ method in the software PopART. For this analysis, we removed samples containing significantly more undefined states based on PopART recommendations.

## Results

### Aphaenogaster Ants

#### Discordance between COI and CAD Consensus Trees and Morphology

Consensus trees for COI and CAD displayed significantly different topologies (RF distance = 148, p = 1; see Figure 1). The CAD tree, for instance, had two large sister clades comprising 35 (henceforth the ‘green’ clade) and 53 (henceforth the ‘yellow’ clade) ants respectively, along with three ants that were basal to both clades. By contrast, the COI tree did not have as strong a cladistic structure. Instead, it was characterized by a paraphyletic clade of 30 basal ants (henceforth the ‘red’ clade) along with another 49 ants nested within the basal clade and comprised of sister clades of 11 (henceforth the ‘purple’ clade) and 38 (henceforth the ‘blue’ clade) ants respectively. Overall, bootstrapped branch support was higher in the COI tree. Despite significant discordance between COI and CAD trees overall (see Figure 1), there was nevertheless detectable congruence between the red, purple and blue COI clades and the green and yellow CAD clades. In particular, ants with red clade COI genes were significantly more likely to have green clade CAD genes, while ants with blue clade COI genes were significantly more likely to have yellow clade CAD genes (Fisher’s exact test: *p =* 0.00067, see SI II Figure S2.1).

**Figure 1.**
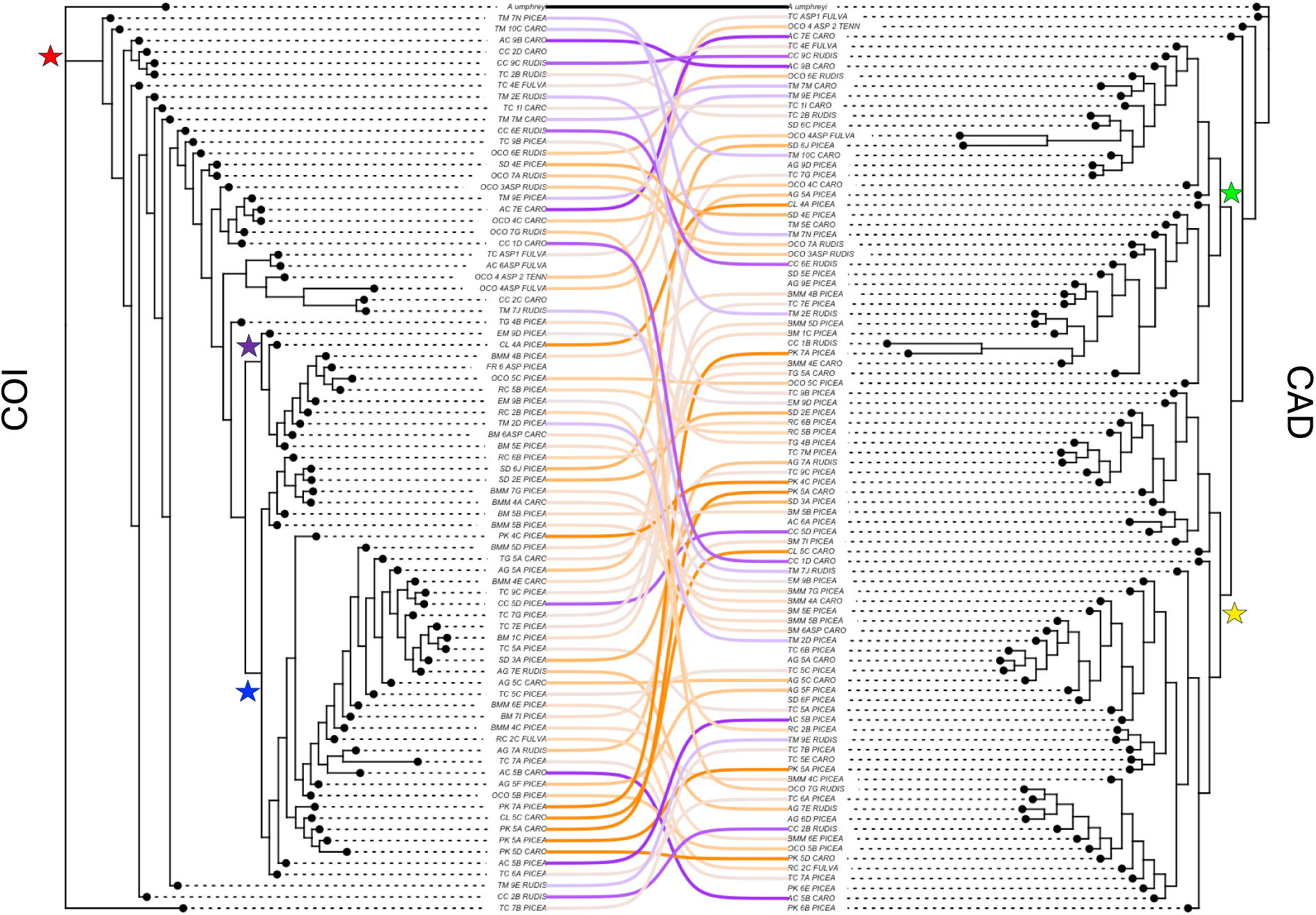
Tanglegram showing high discordance in *A. rudis* complex phylogenetic tree reconstructions based on mitochondrial (left, COI) and nuclear (right, CAD) genes. Colors of the tanglegram lines are based on an east (orange) to west (purple) color scheme. Unpaired samples indicate that sequencing failed for either the COI or CAD gene. The red star marks the node that distinguishes the 30 most basal ants from the outgroup (*A. umphreyi*) in the COI tree, while the blue and purple stars mark the nodes that distinguish the purple and blue clades from both each other and the subtending ‘red’ clade ants. The yellow and green stars mark the nodes that separates the yellow and green clades from each other and from the 3 basal ants and outgroup (*A. umphreyi*) in the CAD tree. Morphological identification is indicated in the sample name (CARO: *A. carolinensis*, FULVA: *A. fulva*, PICEA: *A. picea*, RUDIS: *A. rudis*, TENN: *A. tennesseensis*).

Neither the COI nor the CAD tree aligned well with morphological identification (see Figure 1). There was, however, a significant difference in the relative distributions of morphologically identified *A. rudis*, *A. fulva*, *A. picea* and *A. carolinensis* between the red, purple and blue clades in the COI tree (Fisher’s exact test: *p* = 2.6E-5). Post-hoc testing with a Benjamini-Hochberg correction indicated that this was because *A. picea* was over-represented relative to *A. rudis* and (marginally) *A. carolinensis* in the purple clade as compared to the red clade, and because *A. picea* was over-represented relative to *A. rudis*, *A. carolinensis* and (marginally) *A. fulva* in the blue clade as compared to the red clade. There was no significant difference in the relative distributions of morphologically identified *A. rudis*, *A. fulva*, *A. picea* or *A. carolinensis* between the yellow and green clades in the CAD tree (Fisher’s exact test: *p* = 0.2252).

#### Correlation Among Spatial, Environmental and Host Phylogenetic Distances

Mantel tests based on Spearman (rank), but not Pearson (magnitude) correlation coefficients demonstrated a significant relationship between host relatedness and geographic distance for the COI gene (Spearman: r: 0.1005, p = 0.020; Pearson: r = 0.0703; p = 0.082; Figure 2A). No significant relationships between host relatedness and geographic distance were detected for the CAD gene (Spearman: r: −0.0902, p = 0.978; Pearson: r = −0.0634 ; p = 0.882; Figure 2B). Likewise, Mantel tests based on both Spearman and Pearson correlation coefficients demonstrated a significant relationship between host relatedness and environmental dissimilarity for the COI gene (Spearman: r = 0.0912, p = 0.014; Pearson: r = 0.0685; p = 0.042; Figure 2C) but not for the CAD gene (Spearman: r = −0.0319, p =0.814 ; Pearson: r = −0.0379, p = 0.832; Figure 2D). Distance-based redundancy analysis (dbRDA) and variance partitioning supported results from individual Mantel tests, suggesting that both distance and distance+environment explained relatedness of the COI gene but not of the CAD gene (see SI II Figures S2.2 and S2.3). Because the COI tree better correlated with environmental and geographic features, because *Wolbachia* (see below) is known to impact COI inheritance, and because bootstrap values for nodes were much higher for the COI tree versus the CAD tree, throughout the remainder of the main text, we present results using the COI tree. Comparable results for the CAD tree can be found in Supplemental Information.

**Figure 2.**
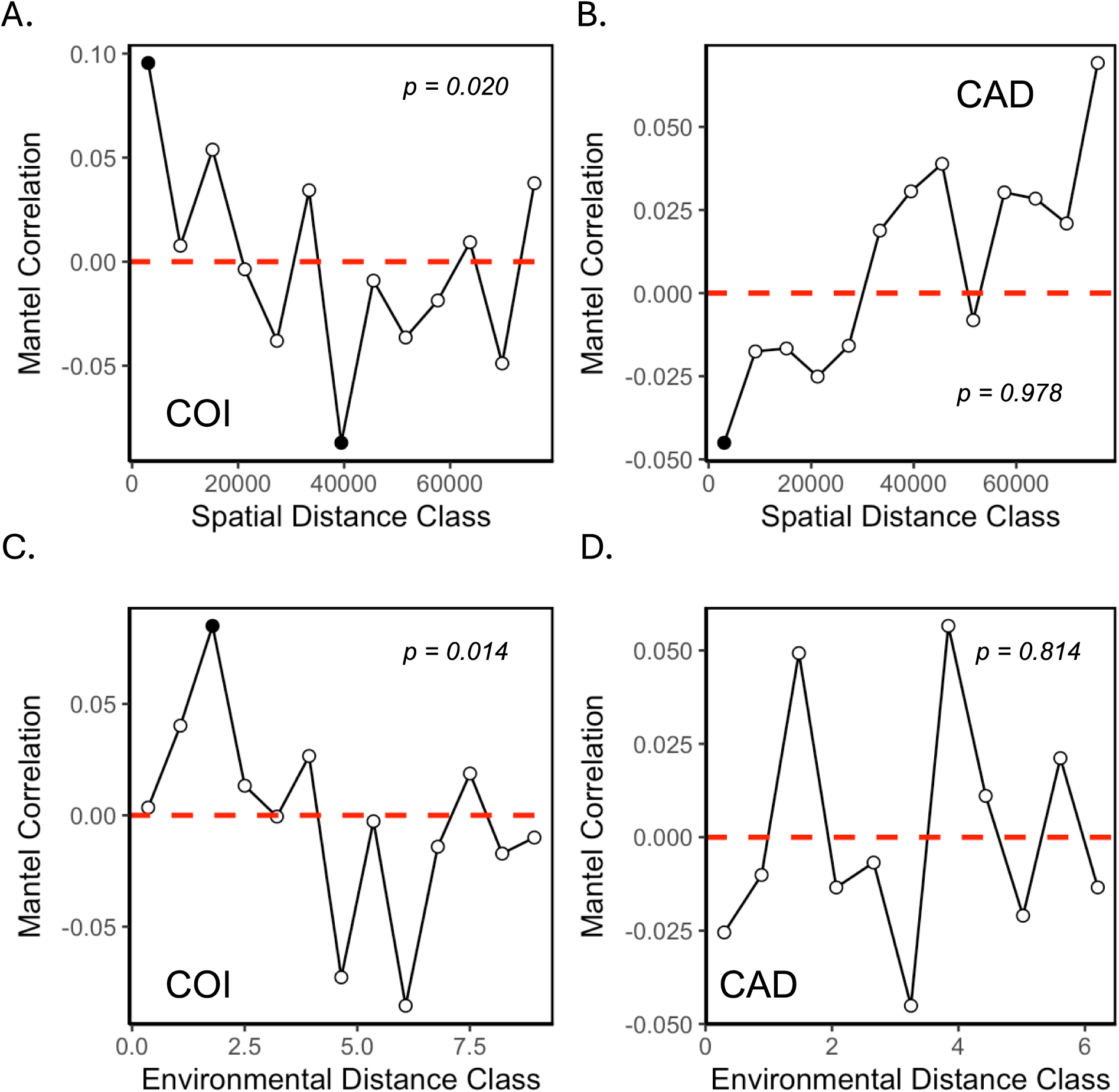
Mantel correlograms for genetic distance based on the COI (A,C) and CAD (B,D) consensus trees, regressed against spatial distance (A,B) and environmental distance (C,D) and assessed with Spearman’s rank correlation. *p-*values shown on each panel reflect the significance of the corresponding Mantel tests (see also SI II Figures S2.2 and S2.3).

#### Differing Spatial and Environmental Distributions of Host Clades

Consistent with the relationships between spatial distance, environmental distance and phylogenetic distance, ants in the red, purple and blue COI clades exhibited different distributions across GSMNP. Red clade ants dominated lower elevations on the Western side of the park, while blue clade ants dominated higher elevations on the Eastern side of the park. Purple clade ants were predominantly found towards the center of the park where they co-occurred with the other two clades across a range of elevations (see Figure 3). In accordance with their differing distributions, environmental conditions were also significantly different across the three COI clades (see Figure 4; Kruskal-Wallis chi-squared = 26.681, df = 2, p-value = 1.6E-6). Ants from the red clade were associated with warmer, drier, and lower elevation conditions while ants from the blue clade were associated with colder, moister and higher elevation conditions. Ants from the purple clade were intermediate, though differences between the blue and the purple clades were not significant. Ancestral state reconstructions for environmental conditions showed similar trends, with more basal ancestral nodes (red clade) being associated with warmer, drier and lower elevation conditions and more derived nodes (purple and blue clades) being associated with cooler, moister and higher elevation conditions (see Figure 4).

**Figure 3.**
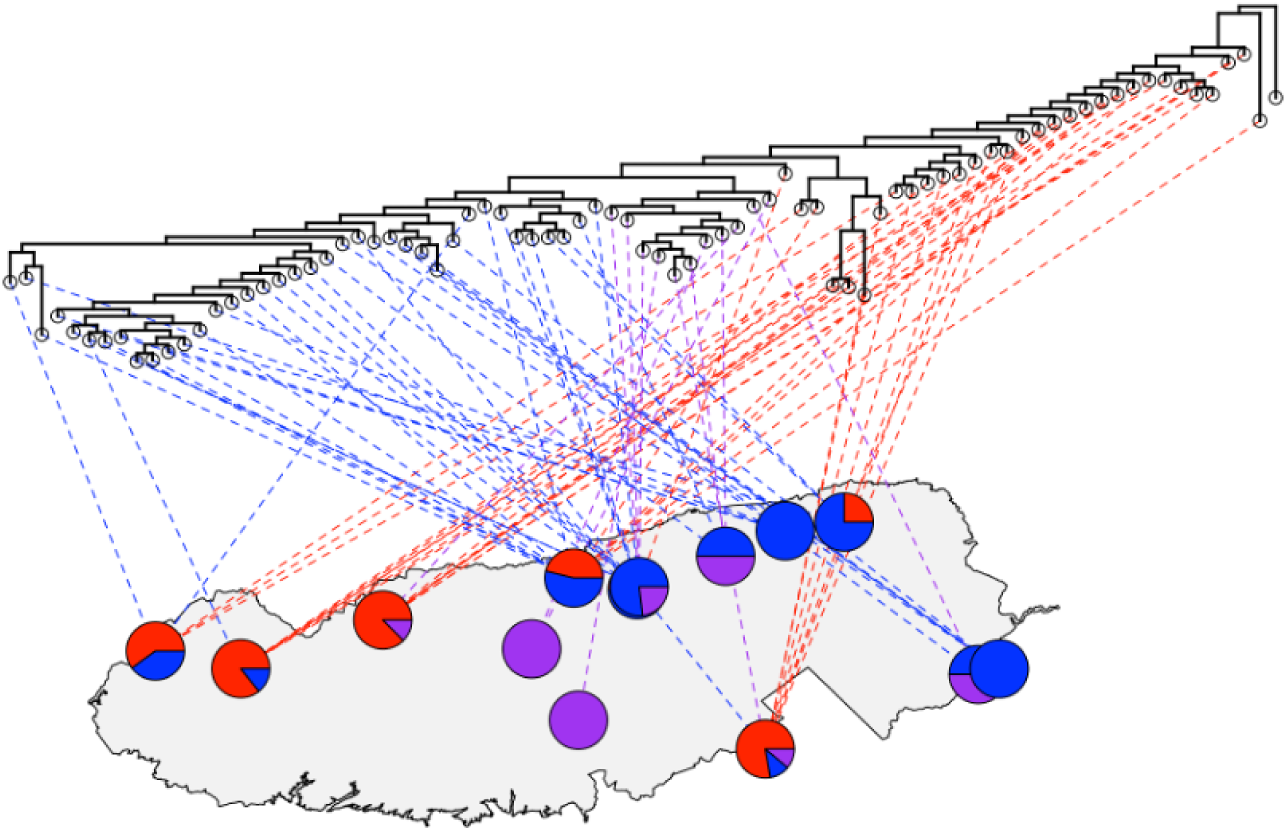
COI phylogenetic tree mapped to 14 sites of origin across 80 ants in GSMNP using the *plot.phylo.to.map* function in the phytools package. The node unassociated with a GSMNP location is the outgroup, *A. umphreyi*. Lines are colored based on whether the ant is included in the red clade or the blue or purple clades. Pie charts show the fraction of ants at each location that come from each clade (see SI II Figure S2.5 for the CAD tree).

**Figure 4.**
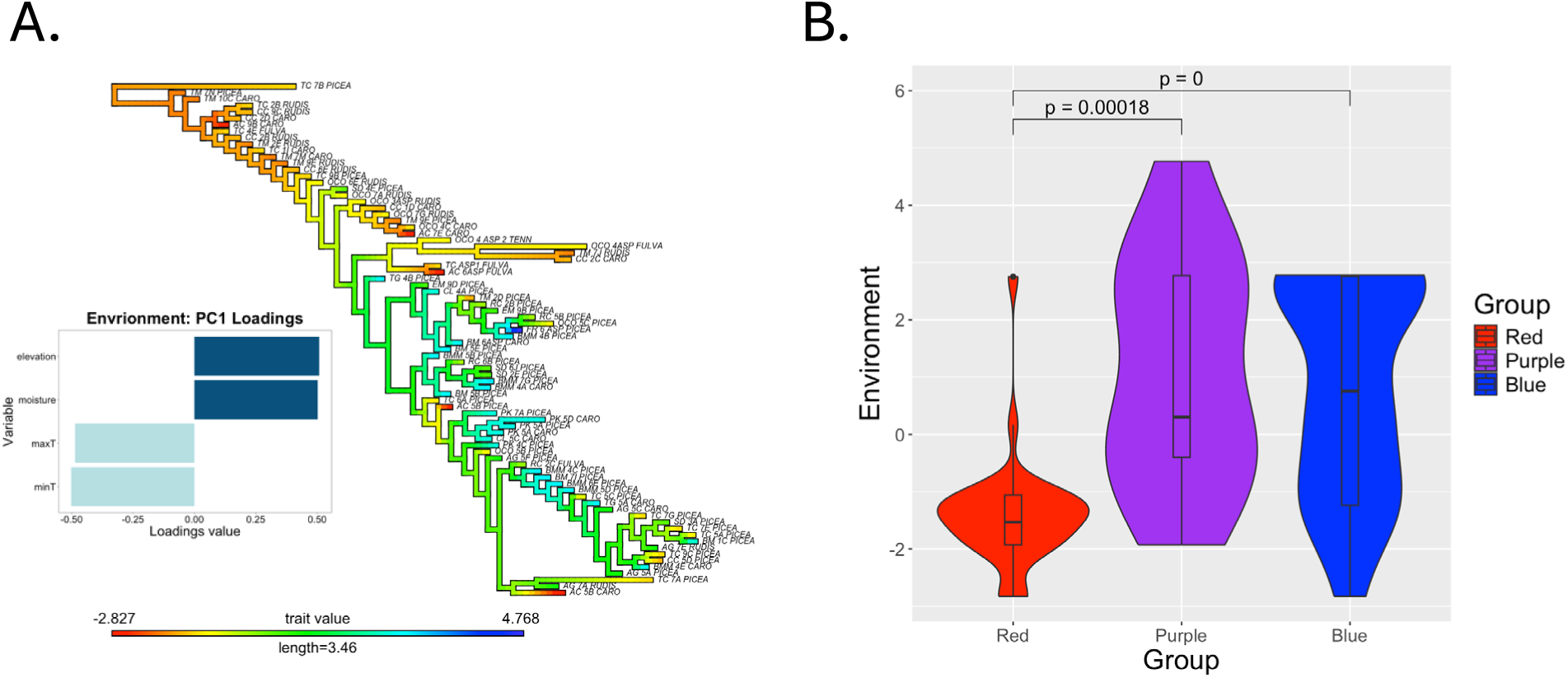
(A) Environmental ancestral trait reconstruction on the COI tree based on a single ‘conglomerate’ environmental axis derived from a PCA analysis on our four environmental variables (minimum soil temperature, maximum soil temperature, soil moisture and elevation). Loadings of the four environmental variable onto the conglomerate environmental axis are shown to the left of the phylogenetic tree. **(B)** Violin plot showing comparison of values along the conglomerate environmental axis for red clade, blue clade and purple clade ants (see SI II Figure S2.6 for comparable results using the CAD tree).

### Aphaenogaster Microbiomes

*A. rudis* complex ant microbiota (see Figure 5) were dominated by Proteobacteria (51% with endosymbionts, 45% without), Firmicutes (22% with endosymbionts, 18% without) and Actinobacteria (9.4% with endosymbionts, 14% without). While some ant microbiota were comprised of a diverse combination of microbial taxa, other profiles were almost entirely comprised of one or two endosymbionts. This was particularly noticeable for *Wolbachia*, although *Spiroplasma* and *Entomoplasma* dominated the microbiota of a few individuals.

**Figure 5.**
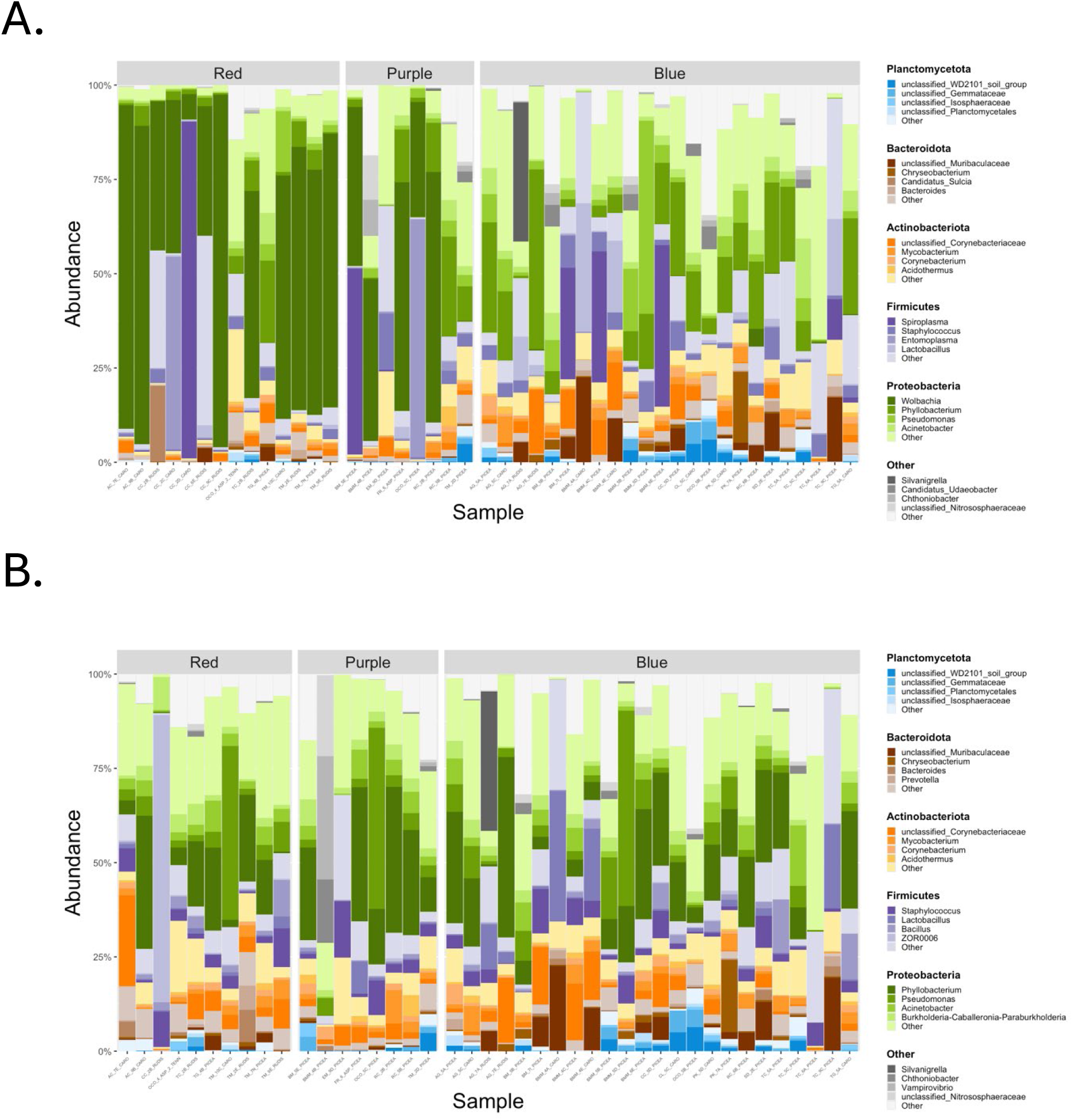
Microbial taxon barplots for *A. rudis* complex ant microbiota with (A) and without (B) dominant endosymbiont genera (*Wolbachia*, *Spiroplasma*, *Entomoplasma*, *Sulcia*). Samples are separated according to COI host clade and colored based on dominant phyla (Planctomycetota: blue, Bacteroidota: brown, Actinobacteriota: orange, Firmicutes: purple, Proteobacteria: green, Other: grey) and genera. Note that several samples from the red clade were dropped due to low reads following removal of endosymbionts (see SI III Figure SI 2.1 for similar barplots divided according to clades within the CAD gene tree).

#### Differential Abundance of Wolbachia, but not Other Endosymbionts, Across Host Clades and Environments

*Wolbachia* relative abundance differed across ant clades, being lowest in the blue clade, intermediate in the purple clade and highest in red clade (Figure 6A; Kruskal-Wallis chi-squared = 26.669, df = 2, p-value = 1.6E-6). Ants from the red clade exhibited a median *Wolbachia* relative abundance of 60.1%, compared to just 0.06% in blue clade ants. Meanwhile, ants from the purple clade exhibited an intermediate relative abundance of 36.7%. Notably, this same pattern held, even when we restricted our analysis to a subset of seven sites where red+purple clade ants co-occurred with blue clade ants (see SI III Figure S3.2B). Thus the lower relative abundance of *Wolbachia* in the blue clade was not entirely a result of spatial and/or environmental effects.

**Figure 6.** (A) Violin plot of *Wolbachia* relative abundance across the three COI ant clades, showing significantly lower abundance in the blue clade. (B) Phylogeny of the 32 *Wolbachia* ASVs in our study, rooted with an uncultured Rickettsiaceae ASV as the outgroup (black). *Wolbachia* tips are colored according to their overall abundance across all ant samples (red: high, white: low). Circular tips are used for non-focal ASVs and diamond tips are used for focal ASVs. Note that the color scheme is logarithmic; the total reads assigned to the three focal strains were wArudF1: 40,033, wArudA1: 21,543 and wArudA2: 186,262. The total reads assigned to the next most abundant *Wolbachia* ASV was 153. (See SI III Figure S3.2 for *Wolbachia* abundances across green and yellow clade ants). (C) Heatmap showing the number of reads of each *Wolbachia* ASV on each ant, organized according to host and microbe phylogenies. Again, note that he color scheme is logarithmic.

Closer inspection suggested that three distinct *Wolbachia* ASVs were responsible for the majority of *Wolbachia* observed in our system (see Figure 6B). Two of the three dominant ASVs (henceforth wArudA1, wArudA2) were highly related, clustering together in supergroup A (see SI I Figure S1.1) and differing from each other by 1/214 base pairs (bp). By contrast, the third dominant ASV (henceforth wArudF1) was highly distinct, clustering in supergroup F and differing from wArudA1 and wArudA2 by 8/214 and 9/214 bp respectively. Notably, these three ASVs showed strong variation across COI ant clades, suggesting a link between mitotype and the presence of specific *Wolbachia* variants (see Figure 6C). ASRs for each of the three *Wolbachia* ASVs provided insight into these patterns (see Figure 7A-C, note that we used ASRs to identify times of strain acquisition/loss, and do not mean to imply any form of *Wolbachia* evolution across the *Aphaenogaster* phylogeny). In particular, ASRs suggested that the ancestral state (red clade) was a microbiota containing all three *Wolbachia* ASVs. wArudF1 and wArudA1, however, were both lost slightly before divergence between the blue and purple clades. Meanwhile, wArudA2 was lost from the blue clade but retained by the purple clade around the time of their divergence. One important caveat is that, although ants from the red, purple and blue clades almost always cluster separately, the precise form of the COI tree is somewhat sensitive to the details of tree construction. More specifically, the red clade is not always ancestral to the blue and purple clades. The haplotype map (see Figure 7D) provides some clarity, suggesting that there are, in fact, three haplotype clusters, with one cluster (the red clade) colonized by all three *Wolbachia* ASVs, one cluster (the purple clade) colonized by wArudA2 and one cluster (the blue clade) having no *Wolbachia*. As expected, the presence/absence of *Wolbachia* ASVs was not strongly conserved within CAD lineages. Nevertheless, the yellow clade had significantly lower *Wolbachia* abundance overall relative to the green clade (see SI III, Figure S3.2A). Consistent with the differing environmental preferences of red, purple and blue clade ants, *Wolbachia* was less abundant in colder, moister, higher elevation regions (Figure 8A; z = −3.242, df = 3, p = 0.00119). Notably, this relationship was significant or marginally significant even when restricting our analysis to only red clade ants (z = −2.139, df = 3, p = 0.0324; Figure 8B) or only blue clade ants (z= −2.237, df = 3 p = 0.0253, Figure 8C when the outlier from Ramsey Cascades was removed, z = −1.818, df = 3, p = 0.0691 otherwise; see SI III Figure S3.6).

**Figure 7.**
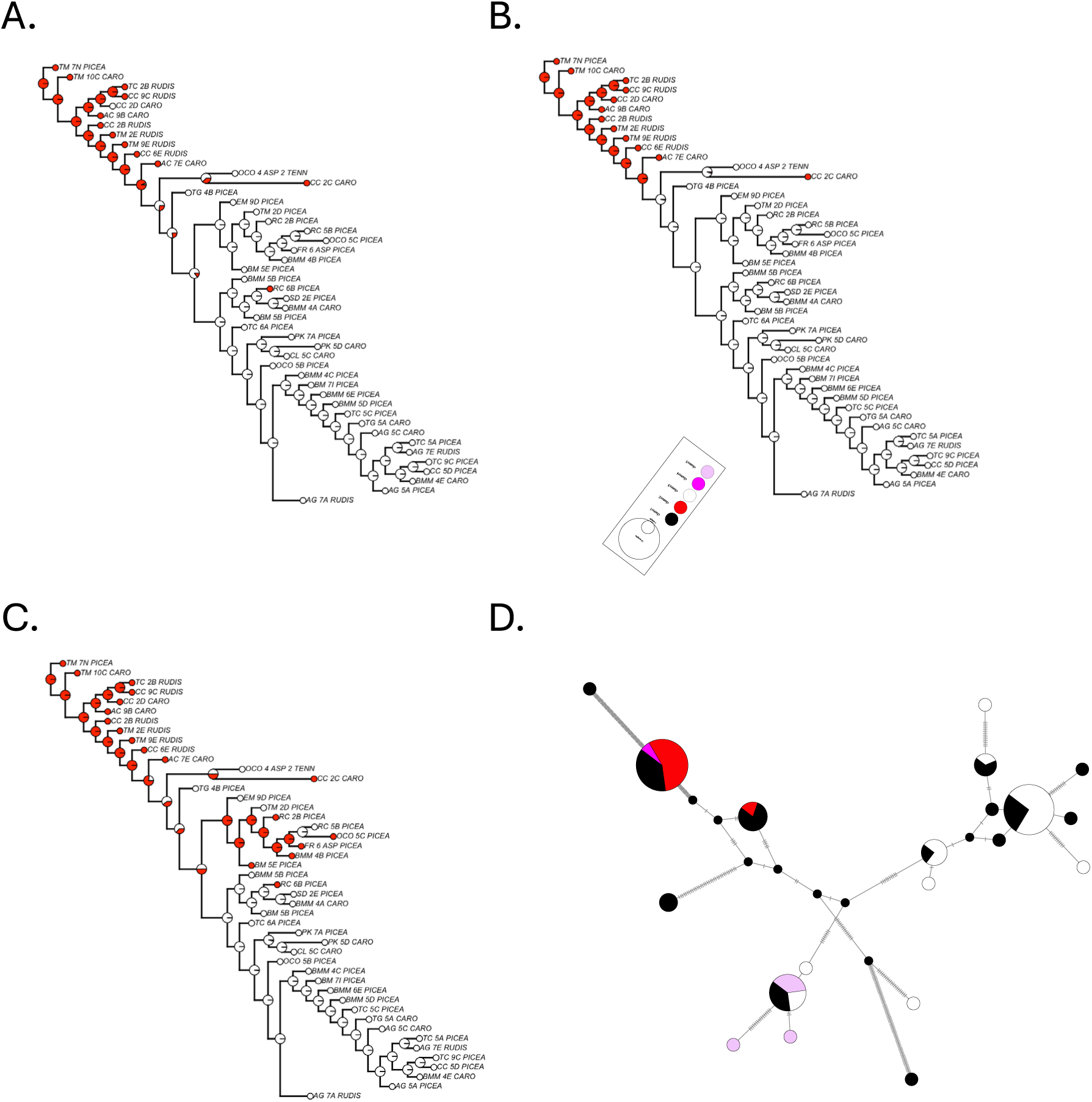
Ancestral state reconstructions for *Wolbachia* wArudF1 **(A)**, wArudA1 **(B)** and wArudA2 **(C)** based on the COI gene tree. **(D)** Haplotype map for COI showing carriage of the three dominant *Wolbachia* ASVs: all three *Wolbachia* ASVs (red), wArudF1 and wArudA1 (dark pink), only wArudA2 (light pink), no *Wolbachia* (white). Black is used for animals for which we had COI sequences but where the HA microbiota did not successfully extract and/or amplify. (See SI III Figure S3.3 for *Wolbachia* ASRs and a haplotype map for the CAD gene tree).

**Figure 8.**
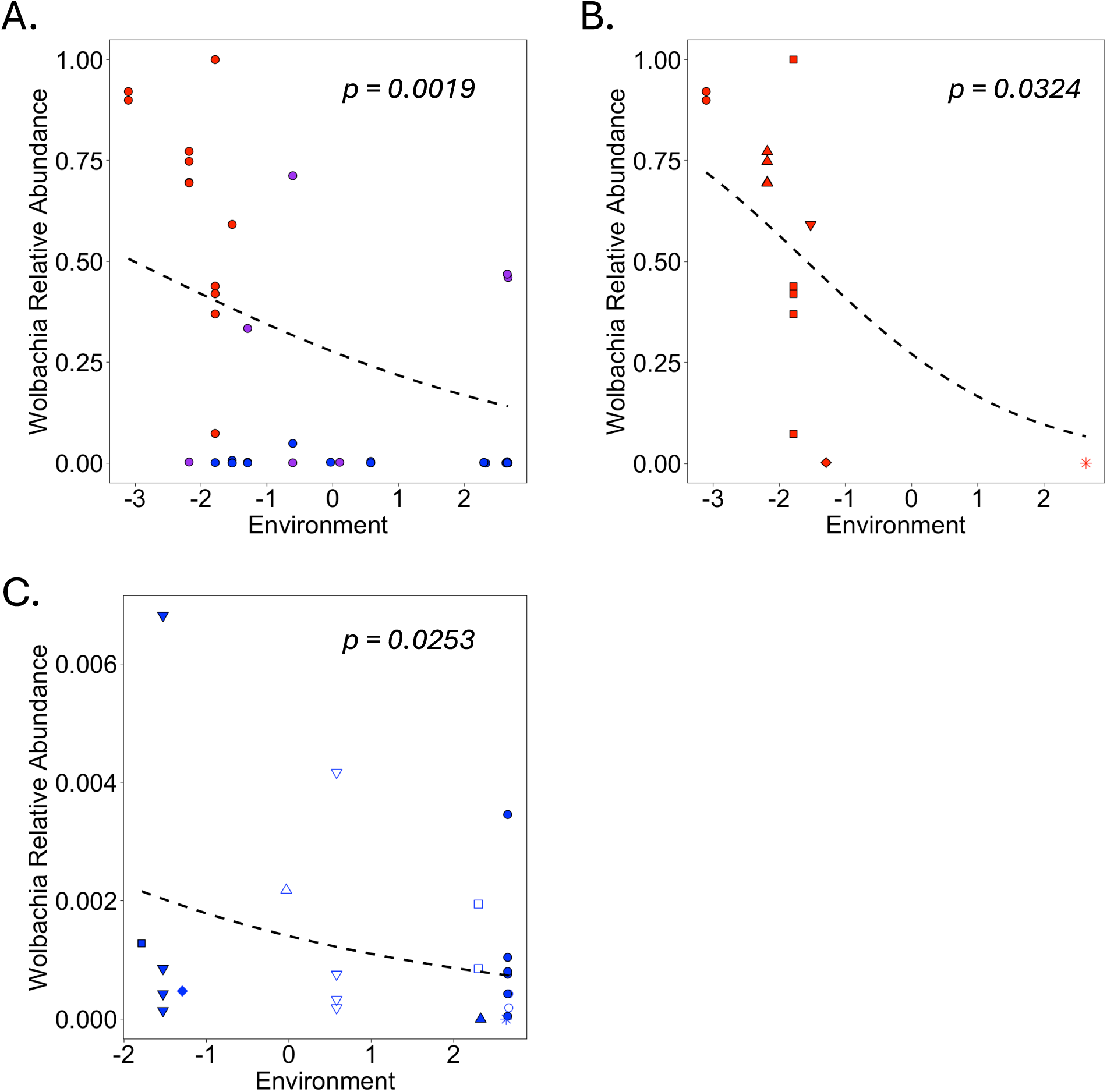
Generalized linear models of *Wolbachia* relative abundance as a function of our conglomerate environmental variable (see Methods) for (A) all ants (B) red clade ants and (C) blue clade ants. COI clades are indicated by the color of the points. In panels (B,C) site is denoted as follows: Abram’s Creek (closed red circles), Tremont (closed upward red triangles), Cades Cove (closed squares), Twin Creeks (closed downward triangles), Oconaluftee (closed diamond), Trillium Gap (star), Snake Den (open upward triangle), Albright Grove (open downward triangle), Cataloochee (closed upward blue triangles), Purchase Knob (open squares), Brush Mountain Myrtle (closed blue circles), Brush Mountain (open circles).

Unlike *Wolbachia*, none of the other endosymbionts that we considered showed differential abundances across ant clades (see SI III Figure 2.2C-E). Further, there was no obvious conservation of these other endosymbionts within specific ant lineages (see SI III Figures S3.4 and S3.5). Rather, almost all these endosymbionts occurred sporadically on a small handful of ants randomly distributed across both space and the COI phylogeny.

#### Non-endosymbiont Microbiota

With endosymbionts removed, Mantel tests failed to detect significant correlations between HA microbiota similarity and host phylogenetic distance based on either the COI or CAD genes (Figure 9A, see also SI III Figures S3.8 and S3.9). There were also no significant correlations between HA microbiota similarity and spatial distance or environmental distance, at least when considering the full set of HA microbiota from all ants (Figure 9B,C). When considering the red and blue clades separately, some metrics (Jaccard, weighted UniFrac) suggested that HA microbiota dissimilarity may increase with increasing spatial distance in the red clade, while other metrics (Jaccard, UniFrac) suggested that HA microbiota dissimilarity may increase with increasing environmental distance in the blue clade (see SI III Figures S3.10-S3.13). Partial Mantel tests and dbRDA suggested a similar lack of variation of HA microbiota with spatial and environmental distance with few and likely spurious exceptions (see SI III Table S3.1). Finally, comparing ants that were positive versus negative for *Wolbachia* indicated minimal differences in HA microbiota composition (Figure 9D), although PERMANOVA was significant or borderline significant for several dissimilarity metrics (Jaccard, UniFrac).

**Figure 9.**
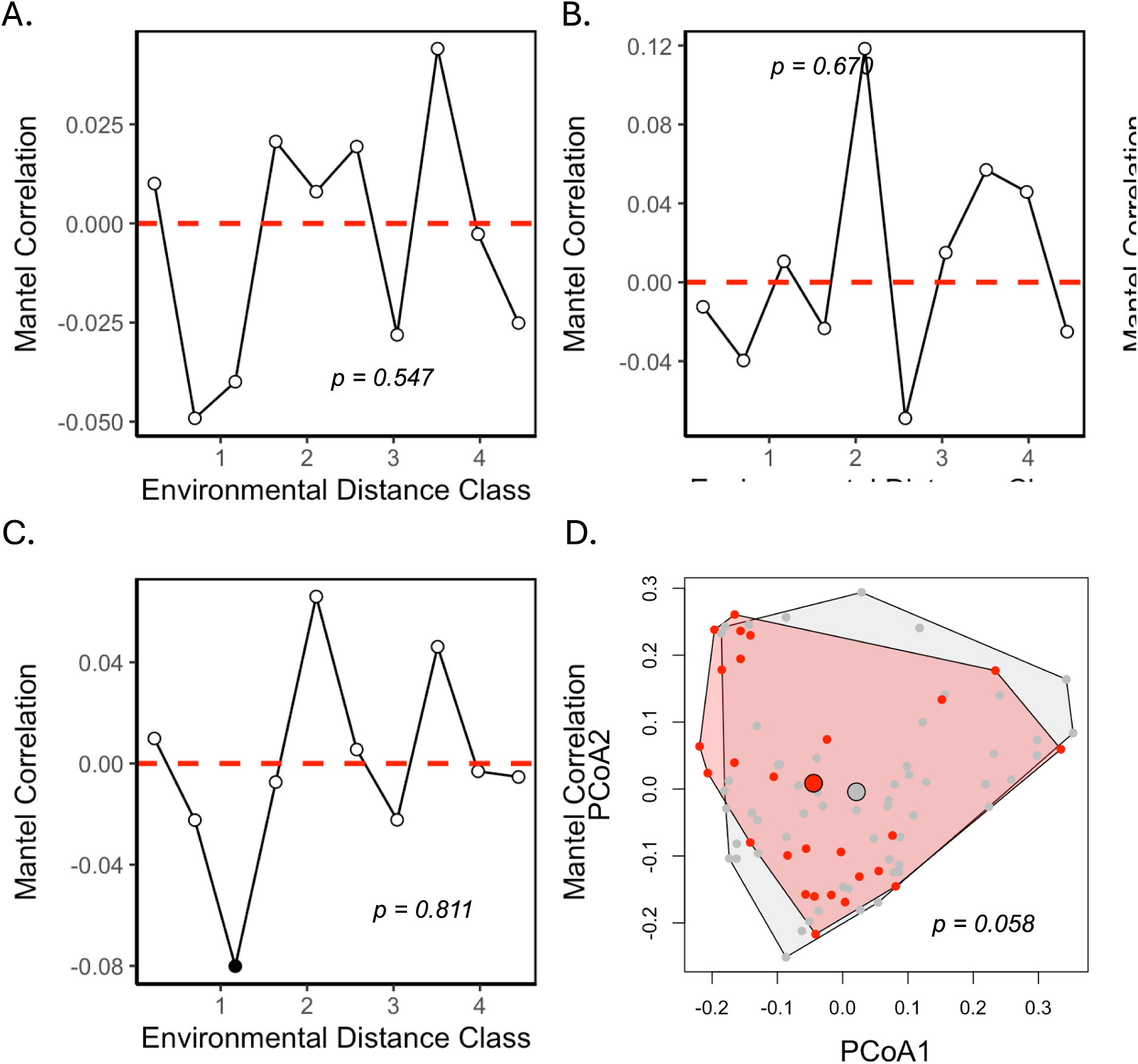
Mantel correlograms for HA microbiota dissimilarity (endosymbionts removed, UniFrac) regressed against host genetic distance (A), spatial distance (B) and environmental distance (C). (D) PCoA (UniFrac) showing the microbiota of hosts that were positive (red) versus negative (grey) for *Wolbachia* prior to removal of endosymbionts. *p*-values on each panel represent the significance of Mantel tests (A-C) and PERMANOVA (D) respectively.

## Discussion

In this paper, we examined the effects of host and microbe scale environmental selection and dispersal limitation on the internal microbiota of *A. rudis* complex ants in Great Smoky Mountains National Park (GSMNP). Interestingly, we found evidence of spatial patterns in both host genetics and the HA microbiota. However, spatial patterns in the HA microbiota were primarily limited to *Wolbachia*. By contrast, the remainder of the HA microbiota, including both other endosymbionts and non-endosymbionts, showed limited effects of either host or microbe scale environmental selection or dispersal limitation.

### Host Scale Environmental Selection and/or Dispersal Limitation impacts Host Genetics

In agreement with previous studies,^39^ we found a significant relationship between host genetics and environment. Interestingly, however, this relationship was only true for COI and not for CAD (Figure 4.2B and Figure 4.2D). For COI gene, we also observed a significant relationship between host genetics and geographic distance (Figure 4.2A and Figure 4.2C). Again, however, this relationship did not hold for CAD. Unfortunately, due to the nature of the ATBI sites^67,68^ and, indeed, the distribution of environments across GSMNP in general, spatial and environmental distances were somewhat confounded in our study. This made it difficult to tease apart the separate effects of host dispersal limitation from host environmental selection. Nevertheless, variance partitioning suggested that both distance and distance + environment caused spatial structure in COI (see SI II Figure S2.3). Thus, both dispersal limitation and environmental selection likely restrict red clade ants to the warmer, drier, lower elevations on the Western side of the park, and blue clade ants to the colder, wetter, higher elevations on the Eastern side of the park.

While the observed spatial structure in COI is interesting, the difference in spatial structure between COI and CAD is equally interesting. This could be driven by differences in either host scale environmental selection or host scale dispersal limitation. In order for differences between COI and CAD to emerge due to host scale environmental selection, the mitochondrial genome must be differentially beneficial in certain environments (i.e., over certain temperature, moisture and/or elevation ranges), while the CAD gene must be largely neutral. This is consistent with the mitochondrial climatic adaptation hypothesis^69^ - a theory that emphasizes the role of mitochondria in determining thermal performance curves (TPCs) and/or critical thermal limits (CTLs).^70^ By contrast, iuf differences between COI and CAD emerge due to host scale dispersal limitation, it implies that mtDNA is more dispersal limited than nuDNA. This is expected purely based on effective population size but could also be exacerbated if female alates are more dispersal limited than males (i.e., sex-biased dispersal).^71^ Notably, higher male dispersal has been documented in other ant species,^72^ though this has not been studied in *Aphaenogaster*.

### Microbe Scale Environmental Selection and/or Dispersal Limitation Impacts HA Microbiota

*Wolbachia* relative abundances significantly differed between COI clades, with red clade individuals containing immensely greater mean relative abundances than blue clade individuals (60.1%% vs 0.06%, respectively). This pattern could emerge either by environmental selection or by a combination of stochastic events and dispersal limitation at either the host or microbe scale. For example, *Wolbachia* may preferentially infect red clade ants because of a shared (but independent) requirement for warmer, drier climates (i.e., host scale environmental selection). Alternatively, *Wolbachia* might preferentially infect red clade ants simply because *Wolbachia* was first introduced into the West side of the park, and barriers to ant movement (i.e., host scale dispersal limitation) have thus far prevented its translocation to blue clade ants on the East side of the park. While both host scale environmental selection and host scale dispersal limitation are plausible, we do not think that either are an adequate explanation for *Wolbachia* distribution in our system. This is because, for either host scale mechanism, differences in *Wolbachia* abundance should only be apparent across environments and/or geographic distance. We found, however, that *Wolbachia* abundances were lower in blue clade ants *even after* controlling for spatial and environmental effects (i.e., in comparisons at sites where both clades co-occur, see Figure S3.2B). This suggests that microbe scale processes are at least partially responsible for observed variation in *Wolbachia* across our system.

Either microbe scale environmental selection or microbe scale dispersal limitation could explain the *Wolbachia* distributions that we found. Red clade ants may, for instance, be more permissive to *Wolbachia* colonization and/or less likely to clear *Wolbachia* colonization once acquired (i.e., microbe scale environmental selection). This would not be unprecedented. Among leaf-cutter ants (*Acromyrmex echinatior*), different patrilines exhibit significant differences in *Wolbachia* densities.^73^ Meanwhile among *Vollenhovia emeryi*, genotypes associated with different wing morphology exhibit different *Wolbachia* colonization rates.^74^ Even among *Aphaenogaster*, a recent study identified differences in *Wolbachia* abundances between *A. rudis* and *A. fulva* ants, although these results were based on only three *A. fulva* and two *A. rudis* colonies.^75^

An alternate explanation is that *Wolbachia* are limited to red clade ants because of microbe scale dispersal limitation. Broadly speaking, *Wolbachia* has two modes of dispersal: horizontal transmission and vertical transmission. Although extensive evidence exists suggesting that *Wolbachia* can be acquired horizontally across large host phylogenetic distances,^76^ including among ants,^77^ horizontal transmission within species requires a number of relatively restrictive conditions.^78^ This likely limits the extent to which *Wolbachia* can transfer directly from one ant to another (microbe scale dispersal limitation), particularly over the shorter, ecological timescales. Consequently, in our system, microbe scale dispersal of *Wolbachia* likely occurs predominantly via vertical transmission (mother to offspring).^21^ As a result, *Wolbachia* colonization should closely mirror patterns in mtDNA inheritance, even if there is no direct effect of the mitochondrial genome on *Wolbachia* colonization rates or abundances (i.e., no microbe scale environmental selection). Again, in our system, it is difficult to determine whether microbe scale environmental selection or dispersal limitation shapes *Wolbachia* carriage. Nevertheless, the extremely tight coupling between COI (i.e., *Aphaenogaster* mitotype) and each of the three *Wolbachia* ASVs, along with the limited number of *Wolbachia* acquisitions and losses predicted by ASRs, suggests that dispersal is strongly limited to vertical transmission. This makes microbe scale dispersal limitation a strong contender for explaining preferential colonization of red clade ants and, by extension, the restriction of *Wolbachia* to the warmer, drier, low elevation regions on the West side of the park.

Unlike *Wolbachia*, we found no evidence of differential abundances between red, purple and/or blue clade ants and any of the other three endosymbionts that we considered. This apparent lack of host effects suggests a corresponding lack of microbe scale environmental selection on *Spiroplasma*, *Entomoplasma* and *Sulcia*. Further, while the generally low prevalence of these other three endosymbionts suggests a role for microbe scale dispersal limitation, this does not seem to lead to emergent spatial patterns, at least at the spatial scales that we considered (ranging from within a site to across the entirety of GSMNP). Indeed, some *Spiroplasma* endosymbionts, while being rare in general, were nevertheless found in all three COI clades and at locations ranging from Abram’s Creek (the lowest elevation Westernmost location) to Cataloochee (a high elevation Easternmost location, see Figure S3.4C). Exactly why *Spiroplasma*, *Entomoplasma* and *Sulcia* exhibit such sporadic distributions relative to *Wolbachia* is unclear. Unraveling differences in the spatial and host occurrences of *Wolbachia* versus other endosybmionts like *Spiroplasma* could shed light on how microbe scale dispersal limitation interacts with other spatial processes to govern the spatial distributions across HA microbial taxa. Similar to non-*Wolbachia* endosymbionts, ANCOM-BC did not find evidence for differential abundances of any other HA microbial taxa across COI clades, nor did PERMANOVA identify compositional differences in general. Again, this suggests that microbe scale environmental selection and/or microbe scale dispersal limitation is less important for microbes other than *Wolbachia*.

### Host Scale Environmental Selection and/or Dispersal Limitation Affect HA Microbiota

In addition to differences in *Wolbachia* abundance across ant clades, we also found strong variation in *Wolbachia* abundance with environment. Admittedly, most of this variation was driven by the varying abundances of the red, purple and blue clade ants in response to environment (i.e., microbe scale environmental selection and/or dispersal limitation, see Figure 8A). However, even within red clade (Figure 8B) and blue clade (Figure 8C) ants, we found evidence for a decrease in *Wolbachia* abundance in colder, drier, higher elevation regions of the park. Thus, in addition to microbe scale processes, host environment (i.e., host scale environmental selection) appears to play a role in governing the spatial distribution of *Wolbachia* This finding is in keeping with a recent global analysis showing that *Wolbachia* colonization tends to increase with increasing temperature in temperate regions.^79^ While many different mechanisms could explain this outcome, one tantalizing possibility is that temperature impacts rates of vertical transmission (i.e., microbe scale dispersal).^80^ Recently documented in *Drosophila*, this mechanism is of particular interest because it would mean that host and microbe scale processes actually *interact* (i.e., host scale environment impacts microbe scale dispersal) to determine spatial patterns in HA microbiota.

While host scale environmental selection appears to impact spatial patterns in HA microbiota, the effects of host scale dispersal limitation are less clear. As suggested above, the lack of spatial structure in CAD sequences indicates potentially high flux of *A. rudis* complex alates across the park, consistent with our initial hypotheses. However, the spatial structure in COI sequences suggests that female movement may be more restricted. If this is true, then any microbes that can be dispersed by males should be relatively free of host scale dispersal limitation. By contrast, microbes that can only be dispersed by females – e.g., primarily vertically transmitted endosymbionts - may be much more strongly impacted by constraints on host dispersal. Interestingly, this once again invokes interaction between host and microbe scale processes (i.e., host scale dispersal limitation only matters because of microbe scale dispersal limitation). The fact that we only see spatial structure in *Wolbachia*, and not in the remainder of the HA microbiota, supports the hypothesis that host scale dispersal limitation may only matter for microbial taxa that also experience strong microbe scale dispersal limitation (i.e., that only disperse through vertical transmission). That said, differences between vertically and horizontally acquired microbes in *A. rudis* could emerge for other reasons (e.g., selection on COI that prevents vertically transmitted microbes, but not horizontally transmitted microbes, from colonizing certain environments, see discussion above).

### HA Microbiota Link with Spatial Patterns in the Host

Although our focus, thus far, has been on identifying the ways in which host and microbial processes combine to shape spatial patterns in HA microbiota, it is also worth speculating on how HA microbiota can shape spatial patterns in the host (i.e., bottom-up mechanisms). There are several possible bottom-up mechanisms, all involving *Wolbachia*, that might explain the spatial structure that we observe in COI and the fact that we do not see the same spatial structure in CAD. First, and most obvious is reproductive manipulation. If *Wolbachia* is entrenched on the Western side of the park, and if *Wolbachia* induces CI or some other form of reproductive manipulation in *A. rudis* ants, then this could prevent Western invasion of blue clade COI, even as blue clade nuclear genes become incorporated into red clade populations. Interestingly, a higher percentage of red clade COI (30%) are mismatched with yellow clade CAD as compared blue clade COI (21%) that are mismatched with green clade CAD, although mismatch in blue clade COI is still considerable, and may be explained by mismatch that existed prior to *Wolbachia* colonization (akin to incomplete lineage sorting), incomplete CI, or even loss of *Wolbachia* entirely by some red clade ants that dispersed East to colder, wetter, higher elevations. Notably, temperature differences can result in differing levels of both CI penetrance and *Wolbachia* density,^81–83^ providing a mechanism by which CI might serve as a strong barrier to gene flow on the Western side of the park, but less of a barrier on the Eastern side of the park.

Like the restriction of blue clade ants to the Eastern side of the park, bottom-up mechanisms could also explain the restriction of red clade ants to the West side of the park. If, for instance, *Wolbachia* colonization lowers dispersal ability, either by altering movement patterns directly or else by reducing production of alates,^84^ then this could slow the Eastern spread of red clade COI, even if CI gives the red clade COI a competitive advantage that should ultimately enable a mitochondrial selective sweep^85^ across the park. Alternately, *Wolbachia* could prevent red clade ants from penetrating Eastward by lowering surviving in colder, wetter, higher elevation regions of the park. Again, this could restrict the spatial extent of the red clade COI, even if it gains a competitive demographic advantage through CI.

### Conclusions and Future Directions

The complex genetic structure of *A. rudis* ants, their propensity to inhabit a wide range of different environments across large spatial extents, and their complex internal HA microbiota comprising both vertically transmitted endosymbionts (e.g., *Wolbachia*, *Spiroplasma*) and horizontally transmitted extracellular taxa makes *A. rudis* an ideal system for studying the interplay between environmental selection and dispersal limitation at the host and microbe scales. Our findings suggest that this interplay leads to interesting spatial patterns across the mountainous terrain in GSMNP. Patterns appear to arise due to both microbe scale processes (differences in *Wolbachia* abundance based on host genetics), as well as host scale processes (differences in *Wolbachia* abundance based on host environment). Further, because these patterns largely involve *Wolbachia*, a well known reproductive manipulator, there is a possibility that bottom-up mechanisms contribute to spatial patterns as well (i.e., the HA microbiota may alter spatial structure in the host). Consequently, this system offers rich possibility for studying bi-directional interactions between hosts and their HA microbes.

Multiple opportunities exist for extending our current study. It would be interesting, for example, to better characterize HA microbiota differences between red, blue and purple clade ants. Now that we have identified a contact zone between these clades, including sites like Oconaluftee where all three co-occur (see Figure 2), it should be possible to more thoroughly explore differences in HA microbiota while controlling for spatial and environmental effects. Likewise, it would be interesting to undertake a more extensive comparison of both the spatial distribution and HA microbiota of ants with and without COI/CAD mismatch. Again, certain locations (e.g. Twin Creeks, Cades Cove, Snakeden) appear to harbor co-occurring ants with different COI and CAD genes, making a more intensive comparison controlling for both spatial and environmental effects possible. Finally, although geographic distance and environment are correlated within GSMNP, it would be interesting to extend our work to the Southern Appalachians more broadly. Because of the many mountaintops and elevational gradients across this region, a broader spatial sampling would allow us to tease apart the effects of environment versus distance, helping to distinguish between processes like host scale environmental selection and host scale dispersal limitation.

Beyond additional field characterization, another interesting next step is to better understand mechanisms driving the spatial patterns that we observe. Lab studies addressing differences in thermal tolerance based on host genotype and *Wolbachia* carriage could, for example, help to explain why red clade ants do not thrive at higher elevations and/or why blue clade ants do not thrive at lower elevations. Likewise, lab studies examining temperature effects on *Wolbachia* load or even whether certain temperature regimes can facilitate *Wolbachia* clearance – a phenomenon that appears somewhat unique to ants^86^ – could help to explain lack of penetrance of *Wolbachia* into higher elevation regions of the park. Similarly, lab studies examining the extent to which *Wolbachia* induces CI or other forms of reproductive manipulation could shed light on whether *Wolbachia* is responsible for spatial structure of host genetics. Finally, understanding how different *Wolbachia* strains interact, and whether this is impacting either HA microbiota or host spatial patterns could be insightful. Indeed, the patterns of *Wolbachia* carriage that we observe, including nested loss across an environmental gradient, are interesting, and could be a valuable system for understanding not only interactions between different *Wolbachia* strains, but also, how multi-strain colonization drives everything from host environmental tolerance to spatial patterns in host evolution.

As keystone mutualists, *Aphaenogaster* ants have garnered significant attention across various research areas,^31,33,37,44,47,87–90^ including recent explorations into their microbial associations.^75,91,92^ Our study suggests that, even beyond their importance to processes like seed dispersal and pest control,^93^ *A. rudis* complex ants might serve as model systems for understanding the multiscale bidirectional interactions between hosts and their HA microbiota.

## Acknowledgments

We thank Dr. Megan Pitz for her help with creation of Figure 3 and her willingness to tolerate centipedes in her closet. We would also like to thank Nico Malagon, Dr. Tatsuhiro Kato, Daniel Cross, Jackson Lloyd, Drew Kanes, Simon Dunn, and Mr. Paul Super for their help in the collection of ant samples. This research was made possible, in part, with support from the Clemson University Genomics and Bioinformatics Facility, which receives support from the College of Science and two Institutional Development Awards (IDeA) from the National Institute of General Medical Sciences of the National Institutes of Health under grant numbers P20GM146584 and P20GM139769.

## Data Accessibility and Benefit-Sharing

### Data Accessibility

All data and code (BIOM files, metadata files, and all R code) necessary for the analyses presented in this manuscript are available at https://github.com/dmalago/Apheanogaster_GSMNP_Microbiome. We will upload raw sequence data (*Aphaenogaster* CO1/CAD and microbial 16S rRNA sequences) to the National Center for Biotechnology Information’s SRA upon acceptance.

### Benefit-Sharing

Research results have been shared with Great Smoky Mountains National Park through yearly reports.

## Author Contributions

DAM, BC and SB designed the experiment and performed the fieldwork. DAM, CH, ERG and MC designed and performed sequencing of the host. DAM, BC, SB, MAS, SM and AS designed and performed sequencing of the microbiota. DAM, SAB, SRB and SRB designed and performed *Wolbachia* analysis. All authors contributed to data analysis and writing of the manuscript.

## References

1. Cottenie, K. Integrating environmental and spatial processes in ecological community dynamics. Ecol Lett 8, 1175–1182 (2005).

2. Soininen, J. Spatial structure in ecological communities – a quantitative analysis. Oikos 125, 160–166 (2016).

3. Leibold, M. A. et al. The metacommunity concept: a framework for multi-scale community ecology. Ecol Lett 7, 601–613 (2004).

4. Bellier, E. et al. Distance decay of similarity, effects of environmental noise and ecological heterogeneity among species in the spatio-temporal dynamics of a dispersal-limited community. Ecography 37, 172–182 (2014).

5. Glück, M., Geue, J. C. & Thomassen, H. A. Environmental differences explain subtle yet detectable genetic structure in a widespread pollinator. BMC Ecol Evol 22, 8 (2022).

6. McRae, B. H. ISOLATION BY RESISTANCE. Evolution (N Y) 60, 1551–1561 (2006).

7. Soininen, J., McDonald, R. & Hillebrand, H. The distance decay of similarity in ecological communities. Ecography 30, 3–12 (2007).

8. Baguette, M., Blanchet, S., Legrand, D., Stevens, V. M. & Turlure, C. Individual dispersal, landscape connectivity and ecological networks. Biological Reviews 88, 310–326 (2013).

9. Couch, C. E. & Epps, C. W. Host, Microbiome, and Complex Space: Applying Population and Landscape Genetic Approaches to Gut Microbiome Research in Wild Populations. Journal of Heredity 113, 221–234 (2022).

10. Couch, C. E. et al. Bighorn sheep gut microbiomes associate with genetic and spatial structure across a metapopulation. Sci Rep 10, (2020).

11. Ruuskanen, M. O., Sommeria-Klein, G., Havulinna, A. S., Niiranen, T. J. & Lahti, L. Modelling spatial patterns in host-associated microbial communities. Environ Microbiol 23, 2374–2388 (2021).

12. Bordenstein, S. R. et al. The disciplinary matrix of holobiont biology. Science (1979) 386, 731–732 (2024).

13. Li, G. et al. Host-microbiota interaction helps to explain the bottom-up effects of climate change on a small rodent species. ISME J 14, 1795–1808 (2020).

14. Risely, A., Klaassen, M. & Hoye, B. J. Migratory animals feel the cost of getting sick: A meta-analysis across species. Journal of Animal Ecology 87, 301–314 (2018).

15. Andriolli, F. S. et al. Do zombie ant fungi turn their hosts into light seekers? Behavioral Ecology 30, 609–616 (2019).

16. Vyas, A., Kim, S.-K., Giacomini, N., Boothroyd, J. C. & Sapolsky, R. M. Behavioral changes induced by Toxoplasma infection of rodents are highly specific to aversion of cat odors. Proceedings of the National Academy of Sciences 104, 6442–6447 (2007).

17. Lemoine, M. M., Engl, T. & Kaltenpoth, M. Microbial symbionts expanding or constraining abiotic niche space in insects. Curr Opin Insect Sci 39, 14–20 (2020).

18. Shamjana, U., Vasu, D. A., Hembrom, P. S., Nayak, K. & Grace, T. The role of insect gut microbiota in host fitness, detoxification and nutrient supplementation. Antonie Van Leeuwenhoek 117, 71 (2024).

19. Otti, O. Genitalia-associated microbes in insects. Insect Sci 22, 325–339 (2015).

20. Telschow, A., Hammerstein, P. & Werren, J. H. The Effect of Wolbachia on Genetic Divergence between Populations: Models with Two-Way Migration. Am Nat 160, S54–S66 (2002).

21. Kaur, R. et al. Living in the endosymbiotic world of Wolbachia: A centennial review. Cell Host Microbe 29, 879–893 (2021).

22. Weinert, L. A., Araujo-Jnr, E. V., Ahmed, M. Z. & Welch, J. J. The incidence of bacterial endosymbionts in terrestrial arthropods. Proceedings of the Royal Society B: Biological Sciences 282, 20150249 (2015).

23. Weeks, A. R., Tracy Reynolds, K. & Hoffmann, A. A. Wolbachia dynamics and host effects: what has (and has not) been demonstrated? Trends Ecol Evol 17, 257–262 (2002).

24. Calvitti, M., Marini, F., Desiderio, A., Puggioli, A. & Moretti, R. Wolbachia Density and Cytoplasmic Incompatibility in Aedes albopictus: Concerns with Using Artificial Wolbachia Infection as a Vector Suppression Tool. PLoS One 10, e0121813 (2015).

25. Tortosa, P. et al. Wolbachia Age-Sex-Specific Density in Aedes albopictus: A Host Evolutionary Response to Cytoplasmic Incompatibility? PLoS One 5, e9700 (2010).

26. Jaenike, J., Dyer, K. A., Cornish, C. & Minhas, M. S. Asymmetrical Reinforcement and Wolbachia Infection in Drosophila. PLoS Biol 4, e325 (2006).

27. Cruz, M. A., Magalhães, S., Sucena, É. & Zélé, F. Wolbachia and host intrinsic reproductive barriers contribute additively to postmating isolation in spider mites. Evolution (N Y) 75, 2085–2101 (2021).

28. Fridley, J. D. Downscaling Climate over Complex Terrain: High Finescale (<1000 m) Spatial Variation of Near-Ground Temperatures in a Montane Forested Landscape (Great Smoky Mountains)*. J Appl Meteorol Climatol 48, 1033–1049 (2009).

29. Stark, J. R. & Fridley, J. D. Topographic Drivers of Soil Moisture Across a Large Sensor Network in the Southern Appalachian Mountains (USA). Water Resour Res 59, (2023).

30. Drake, V. A. & Farrow, R. A. The Influence of Atmospheric Structure and Motions on Insect Migration. Annu Rev Entomol 33, 183–210 (1988).

31. Lubertazzi, D. The biology and natural history of Aphaenogaster rudis. Psyche (Stuttg) (2012) doi:10.1155/2012/752815.

32. Paysen, E. E. S. DIVERSITY AND ABUNDANCE OF ANTS AT FOREST EDGES IN THE GREAT SMOKY MOUNTAINS NATIONAL PARK. undefined-undefined (2007).

33. Helms Cahan, S., et al. Modulation of the heat shock response is associated with acclimation to novel temperatures but not adaptation to climatic variation in the ants Aphaenogaster picea and A. rudis. Comp Biochem Physiol A Mol Integr Physiol 204, 113–120 (2017).

34. Lessard, J.-P., Dunn, R. R., Parker, C. R. & Sanders, N. J. Rarity and Diversity in Forest Ant Assemblages of Great Smoky Mountains National Park. Southeastern Naturalist 6, 215–228 (2007).

35. Lynch, J. F. Seasonal, Successional, and Vertical Segregation in a Maryland Ant Community. Oikos 37, 183 (1981).

36. Fellers, J. H. Interference and exploitation in a guild of woodland ants. Ecology 68, 1466–1478 (1987).

37. Banschbach, V. S., Brunelle, A., Bartlett, K. M., Grivetti, J. Y. & Yeamans, R. L. Tool use by the forest ant Aphaenogaster rudis: Ecology and task allocation. Insectes Soc 53, 463–471 (2006).

38. Cole, A. C. A Guide to the Ants of the Great Smoky Mountains National Park, Tennessee. American Midland Naturalist 24, 1–88 (1940).

39. Warren, R. J. et al. Climate-driven range shift prompts species replacement. Insectes Soc 63, 593–601 (2016).

40. Warren, R. J., McAfee, P. & Bahn, V. Ecological differentiation among key plant mutualists from a cryptic ant guild. Insectes Soc 58, 505–512 (2011).

41. Buono, C. M. et al. Uncovering how behavioral variation underlying mutualist partner quality is partitioned within a species complex of keystone seed-dispersing ants. Insectes Soc 69, 247–260 (2022).

42. Nichols, B. J. & Langdon, K. R. The Smokies All Taxa Biodiversity Inventory: History and Progress. Southeastern Naturalist 6, (2007).

43. Ferro, M. A $3 Homemade Winkler Sampler. Newsletter of the International Society of Hymenopterists 9, 5–7 (2018).

44. Umphrey, G. J. Morphometric discrimination among sibling species in the fulva – rudis – texana complex of the ant genus Aphaenogaster (Hymenoptera: Formicidae). Can J Zool 74, 528–559 (1996).

45. Brady, S. G., Schultz, T. R., Fisher, B. L. & Ward, P. S. Evaluating alternative hypotheses for the early evolution and diversification of ants. Proceedings of the National Academy of Sciences 103, 18172–18177 (2006).

46. Ward, P. S., Brady, S. G., Fisher, B. L. & Schultz, T. R. Phylogeny and biogeography of dolichoderine ants: Effects of data partitioning and relict taxa on historical inference. Syst Biol 59, 342–362 (2010).

47. DeMarco, B. B. & Cognato, A. I. A multiple-gene phylogeny reveals polyphyly among eastern North American Aphaenogaster species (Hymenoptera: Formicidae). Zool Scr 45, 512–520 (2016).

48. Caterino, M. S. & Harden, C. W. Unseeing and Unseen: On the Distribution, Morphology, and Larva of One of North America’s Rarest Histerid Beetles, Geocolus caecus Wenzel (Coleoptera: Histeridae). Coleopt Bull 76, (2022).

49. Team, R. C. & R Core Team. R Core Team (2014). R: A language and environment for statistical computing. R Foundation for Statistical Computing, Vienna, Austria. URL http://www.R-project.org/. R Foundation for Statistical Computing (2014).

50. Chao, K.-H., Barton, K., Palmer, S. & Lanfear, R. sangeranalyseR: Simple and Interactive Processing of Sanger Sequencing Data in R. Genome Biol Evol 13, (2021).

51. Trifinopoulos, J., Nguyen, L.-T., von Haeseler, A. & Minh, B. Q. W-IQ-TREE: a fast online phylogenetic tool for maximum likelihood analysis. Nucleic Acids Res 44, W232–W235 (2016).

52. Letten, A. D. & Cornwell, W. K. Trees, branches and (square) roots: why evolutionary relatedness is not linearly related to functional distance. Methods Ecol Evol 6, 439–444 (2015).

53. Kozich, J. J., Westcott, S. L., Baxter, N. T., Highlander, S. K. & Schloss, P. D. Development of a Dual-Index Sequencing Strategy and Curation Pipeline for Analyzing Amplicon Sequence Data on the MiSeq Illumina Sequencing Platform. Appl Environ Microbiol 79, 5112–5120 (2013).

54. Bolyen, E. et al. Reproducible, interactive, scalable and extensible microbiome data science using QIIME 2. Nat Biotechnol 37, 852–857 (2019).

55. Callahan, B. J. et al. DADA2: High-resolution sample inference from Illumina amplicon data. Nat Methods 13, (2016).

56. McMurdie, P. J. & Holmes, S. Phyloseq: An R Package for Reproducible Interactive Analysis and Graphics of Microbiome Census Data. PLoS One 8, (2013).

57. Breeuwer, J. A. & Werren, J. H. Cytoplasmic incompatibility and bacterial density in Nasonia vitripennis. Genetics 135, 565–574 (1993).

58. Faith, D. P. Conservation evaluation and phylogenetic diversity. Biol Conserv 61, 1–10 (1992).

59. Lozupone, C. & Knight, R. UniFrac: a New Phylogenetic Method for Comparing Microbial Communities. Appl Environ Microbiol 71, 8228–8235 (2005).

60. Mantel, N. The detection of disease clustering and a generalized regression approach. Cancer Res (1967).

61. Jari Oksanen, Roeland Kindt, Pierre Legendre, Bob O’Hara, Gavin L. Simpson, P. & Solymos, M. Henry H. Stevens, H. W. Package ‘vegan’. https://www.researchgate.net/publication/228339454_The_vegan_Package?enrichId=rgreq-87dd17328dd562faca4cd6cd67fd8d2b-XXX&enrichSource=Y292ZXJQYWdlOzIyODMzOTQ1NDtBUzoxMDIxMjA3NjYzMTI0NjVAMTQwMTM1ODg5NzY3NQ%3D%3D&el=1_x_2&_esc=publicationCoverPdf http://cran.ism.ac.jp/web/packages/vegan/vegan.pdf (2007).

62. Paradis, E. & Schliep, K. ape 5.0: an environment for modern phylogenetics and evolutionary analyses in {R}. Bioinformatics 35, 526–528 (2019).

63. Jaccard, P. THE DISTRIBUTION OF THE FLORA IN THE ALPINE ZONE. 1. New Phytologist 11, 37–50 (1912).

64. Bray, J. R. & Curtis, J. T. An Ordination of the Upland Forest Communities of Southern Wisconsin. Ecol Monogr 27, 325–349 (1957).

65. Nelder, J. A. & Wedderburn, R. W. M. Generalized Linear Models. J R Stat Soc Ser A 135, 370 (1972).

66. Templeton, A. R., Crandall, K. A. & Sing, C. F. A cladistic analysis of phenotypic associations with haplotypes inferred from restriction endonuclease mapping and DNA sequence data. III. Cladogram estimation. Genetics 132, 619–633 (1992).

67. Jenkins, Michael, A. Vegetation Communities of Great Smoky Mountains National Park. Southeastern Naturalist (2007) 10.1656/1528-7092(2007)6[35:VCOGSM]2.0.CO;2.

68. Kanes, D. et al. Species Distribution Models Reveal Varying Degrees of Refugia From the Invasive Asian Needle Ant for Native Ants Versus Ant-Plant Seed Dispersal Mutualisms. Ecol Evol 15, (2025).

69. Camus, M. F., Wolff, J. N., Sgrò, C. M. & Dowling, D. K. Experimental Support That Natural Selection Has Shaped the Latitudinal Distribution of Mitochondrial Haplotypes in Australian Drosophila melanogaster. Mol Biol Evol 34, 2600–2612 (2017).

70. Chung, D. J. & Schulte, P. M. Mitochondria and the thermal limits of ectotherms. Journal of Experimental Biology 223, (2020).

71. Toews, D. P. L. & Brelsford, A. The biogeography of mitochondrial and nuclear discordance in animals. Mol Ecol 21, 3907–3930 (2012).

72. Sundström, L., Keller, L. & Chapuisat, M. INBREEDING AND SEX-BIASED GENE FLOW IN THE ANT FORMICA EXSECTA. Evolution (N Y) 57, 1552–1561 (2003).

73. Frost, C. L., Mitchell, R., Smith, J. E. & Hughes, W. O. H. Genotypes and phenotypes in a Wolbachia -ant symbiosis. PeerJ 12, e17781 (2024).

74. Noh, P. et al. Association between host wing morphology polymorphism and Wolbachia infection in Vollenhovia emeryi (Hymenoptera: Myrmicinae). Ecol Evol 10, 8827–8837 (2020).

75. Kelleher, L. A. & Ramalho, M. O. Impact of Species and Developmental Stage on the Bacterial Communities of Aphaenogaster Ants. Curr Microbiol 82, 157 (2025).

76. Ahmed, M. Z., Breinholt, J. W. & Kawahara, A. Y. Evidence for common horizontal transmission of Wolbachia among butterflies and moths. BMC Evol Biol 16, 118 (2016).

77. Ramalho, M. de O., Kim, Z., Wang, S. & Moreau, C. S. Wolbachia Across Social Insects: Patterns and Implications. Ann Entomol Soc Am 114, 206–218 (2021).

78. Tolley, S. J. A., Nonacs, P. & Sapountzis, P. Wolbachia Horizontal Transmission Events in Ants: What Do We Know and What Can We Learn? Front Microbiol 10, (2019).

79. Charlesworth, J., Weinert, L. A., Araujo, E. V. & Welch, J. J. Wolbachia, Cardinium and climate: an analysis of global data. Biol Lett 15, 20190273 (2019).

80. Hague, M. T. J. et al. Temperature effects on cellular host-microbe interactions explain continent-wide endosymbiont prevalence. Current Biology 32, 878–888.e8 (2022).

81. Hurst, G. D. D., Johnson, A. P., Schulenburg, J. H. G. v d & Fuyama, Y. Male-Killing Wolbachia in Drosophila: A Temperature-Sensitive Trait With a Threshold Bacterial Density. Genetics 156, 699–709 (2000).

82. Mouton, L., Henri, H., Bouletreau, M. & Vavre, F. Effect of temperature on Wolbachia density and impact on cytoplasmic incompatibility. Parasitology 132, 49–56 (2006).

83. Bordenstein, S. R. & Bordenstein, S. R. Temperature Affects the Tripartite Interactions between Bacteriophage WO, Wolbachia, and Cytoplasmic Incompatibility. PLoS One 6, e29106 (2011).

84. Wenseleers, T., Sundström, L. & Billen, J. Deleterious Wolbachia in the ant Formica truncorum. Proc R Soc Lond B Biol Sci 269, 623–629 (2002).

85. Cariou, M., Duret, L. & Charlat, S. The global impact of Wolbachia on mitochondrial diversity and evolution. J Evol Biol 30, 2204–2210 (2017).

86. Russell, J. A. The ants (Hymenoptera: Formicidae) are unique and enigmatic hosts of prevalent Wolbachia (Alphaproteobacteria) symbionts. Myrmecol News (2011).

87. Nelson, B. Y. INCREASED FUNGAL DIVERSITY ASSOCIATED WITH APHAENOGASTER SPP.: MORE EVIDENCE FOR KEYSTONE MUTUALISMS. (Western Carolina University, 2012).

88. University, M. N. S.-W. C. & undefined 2014. Effects of myrmecochore species abundance, diversity, and fruiting phenology on Aphaenogaster (Hymenoptera: Formicidae) nesting and foraging in Southern. core.ac.ukMN SchultzWestern Carolina University, 2014•core.ac.uk (2014).

89. Ness, J. H., Morin, D. F. & Giladi, I. Uncommon specialization in a mutualism between a temperate herbaceous plant guild and an ant: Are Aphaenogaster ants keystone mutualists? Oikos 118, 1793–1804 (2009).

90. Paysen, E. S. et al. EFFECTS OF MYRMECOCHORE SPECIES ABUNDANCE, DIVERSITY, AND FRUITING PHENOLOGY ON APHAENOGASTER (HYMENOPTERA: FORMICIDAE) NESTING AND FORAGING IN SOUTHERN APPALACHIAN RICH COVE FORESTS. Ecology 10, 5–7 (2018).

91. Lenoir, A. & Devers, S. Alkaloid secretion inhibited by antibiotics in Aphaenogaster ants. C R Biol 341, 358–361 (2018).

92. Pagalilauan, A., Pavloudi, C., Ospina, S. M., Smith, A. & Saw, J. Interaction with refuse piles is associated with co-occurrence of core gut microbiota in workers of the ant Aphaenogaster picea. Preprint at 10.1099/acmi.0.000832.v3 (2025).

93. Perfecto, I. & Philpott, S. M. Ants (Hymenoptera: Formicidae) and ecosystem functions and services in urban areas: a reflection on a diverse literature. Myrmecol News (2023).

94. Boulanger, E. et al. Climate differently influences the genomic patterns of two sympatric marine fish species. Journal of Animal Ecology 91, 1180–1195 (2022).

