## Supplemental Information I for "Host and Microbe Scale Processes Shape Spatial Variation in *Aphaenogaster* (Hymenoptera: Formicidae) Genetics and Their Microbiota"

**Supplemental Information I: Additional Methods**

*Ant Collection*

To collect ants from each quadrat of litter we scraped the leaf litter and top ~1 cm of soil from each quadrat and sifted it using a litter sieve to separate larger debris from fine litter and arthropods. We sifted until no small debris was left in the top aspect of the sifter and then placed all collected fine litter in a cotton bag along with a unique identifier tag. Bags were tied at the top to prevent any arthropods from escaping and were kept moist until they could be returned to our lab for processing (typically 3-5 days). To extract litter arthropods, samples were hung in a temperature-controlled laboratory at 20°C using self-assembled winkler extractors[43] for 12 days. Winkler extractors were shaken 2 days after initial hanging to encourage arthropod movement. Collected ants were stored in 100% ethanol until identification. Ant sampling was conducted in GSMNP under permit number GRSM-2020-SCI-2463.

*Ant Identification*

DAM and BC identified collected ants to the genus level using the key found in Ants of North America: A Guide to the Genera. *A. rudis* complex ants were then identified to species by BC by using a species complex key.[44] Following identification, ants were kept frozen at -80°C in 1.5mL tubes filled with 100% ethanol until they could be characterized genetically through sequencing of the mitochondrial Cytochrome Oxidase I (COI), and nuclear carbomoylphosphate synthase (CAD) genes (see below). These loci contain phylogenetically informative nucleotide positions previously used for *Aphaenogaster* species delimitation and represent one mitochondrial (mtDNA) and one nuclear (nuDNA) marker respectively.[45–47]

*Microbial ASV Taxonomic Assignment and Subsequent Dataset Manipulation*

FASTQ files obtained through sequencing (see Main Text) were analyzed using a custom Qiime2[54] pipeline (see Github repo). We then filtered and processed reads using DADA2[55] with default trim and length settings changed to: --p-trim-left-f 19, --p-trim-left-r 20, --p-trunc-len-f 225, --p-trunc-len-r 225. We grouped sequenced reads into amplicon sequence variants (ASVs). A phylogenetic tree of our sequenced reads was created using the Qiime2 *align-to-tree-mafft-fasttree* function. Next, we trained a naïve bayes classifier specific to our primer set using the *feature-classifier* Qiime2 function. Finally, ASVs were taxonomically classified using the Silva 99% similarity database and outputted as BIOM files for subsequent manipulation in R (version 4.2.1), primarily using functions from the *phyloseq* package (version 1.50.0).[56]

For full microbiota analyses, we first removed all sequences which could not be classified to phylum, as well as all mitochondrial and chloroplast sequences. This left only bacterial and archaeal amplicon sequence variants (ASVs). In addition, we removed potential contaminants by subtracting the maximum number of reads of each ASV found in any of our 19 sequencing blanks. Following this cleaning step, we removed all microbiota samples with a final read depth less than 20,000 reads, considering these sequencing failures. We then rarified to the number of reads in the lowest remaining sample (22,533 reads per sample). This left a total of 62 samples. We used all 62 of these samples for analyses involving only microbiota. For analyses that paired microbiota and COI/CAD, we only used samples for which both host and microbiota data were available (see Main Text).

Because we normalized library size across samples, and because endosymbionts, particularly *Wolbachia* (see Main Text) can dominate samples where they occur, we repeated all of our microbiota analyses removing ASVs mapping to dominant endosymbiotic genera

(*Wolbachia, Spiroplasma*, *Entomoplasma* and *Sulcia*). Endosymbiont reads were removed from the full microbiota dataset prior to rarefication and all microbiota samples were then rarified to a final read depth of 10,000. Again, we used all available microbiota samples for microbiota analyses. For analyses that paired microbiota and COI/CAD, we only used samples for which both host and microbiota data were available.

*Wolbachia Supergroups*

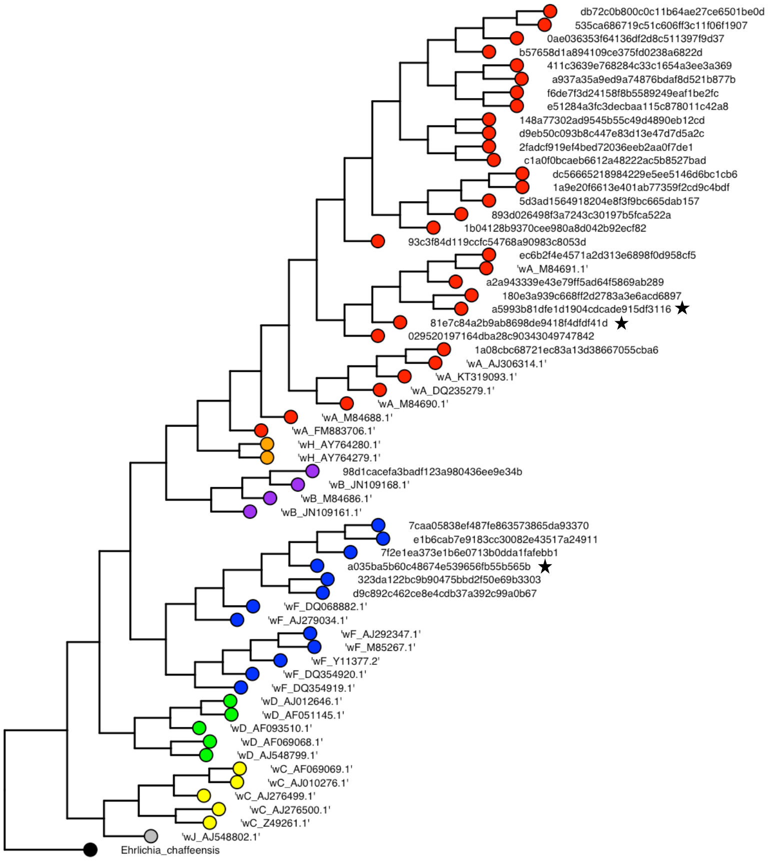


**Figure S1.1** Identification of *Wolbachia* supergroups. Reference *Wolbachia* strains are labelled with a 'w', followed by the supergroup and NCBI accession number. All other strains are from the current study, with the three dominant *Wolbachia* ASVs marked by black stars. *Ehrlichia chaffeensis* is used as an outgroup (black). Supergroups are colored as follows: *A*: red, *B*: purple, *C*: yellow, *D*: green, *F*: blue, *H*: orange, *J*: grey.

### *Endosymbiont Thresholds*

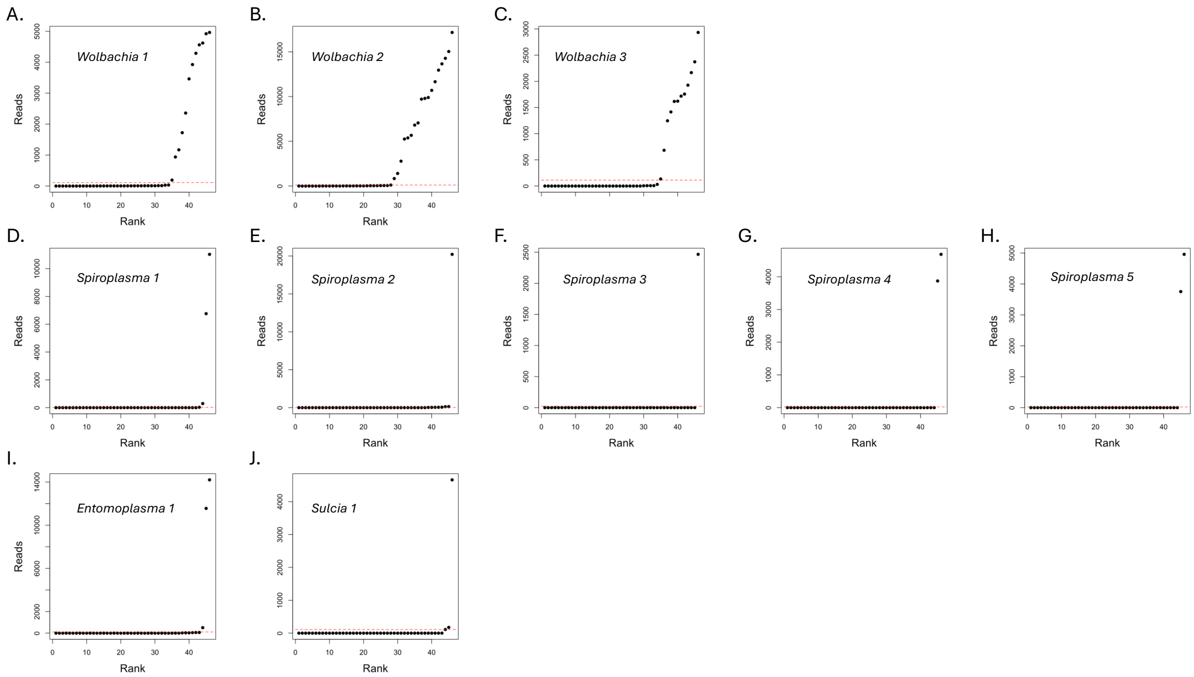


**Figure S1.2** Read counts for each of the three *Wolbachia* (A-C), five *Spiroplamsa* (D-H), one *Entomoplasma* (I) and one *Sulcia* (J) strains, ranked from lowest to highest and with the 0.5% (A-C) and 0.1% (D-J) thresholds marked using a dashed red line.

**Figure S1.2** shows how the thresholds for determining *Wolbachia*, *Spiroplasma*, *Entomoplasma* and *Sulcia* presences versus absences were determined. More specifically, 0.5% marks an ‘elbow’ that is largely consistent across the *Wolbachia* strains, while 0.1% marks an ‘elbow’ that is consistent across the remaining endosymbiont strains.

**Supplemental Information II: Additional Host Analyses**

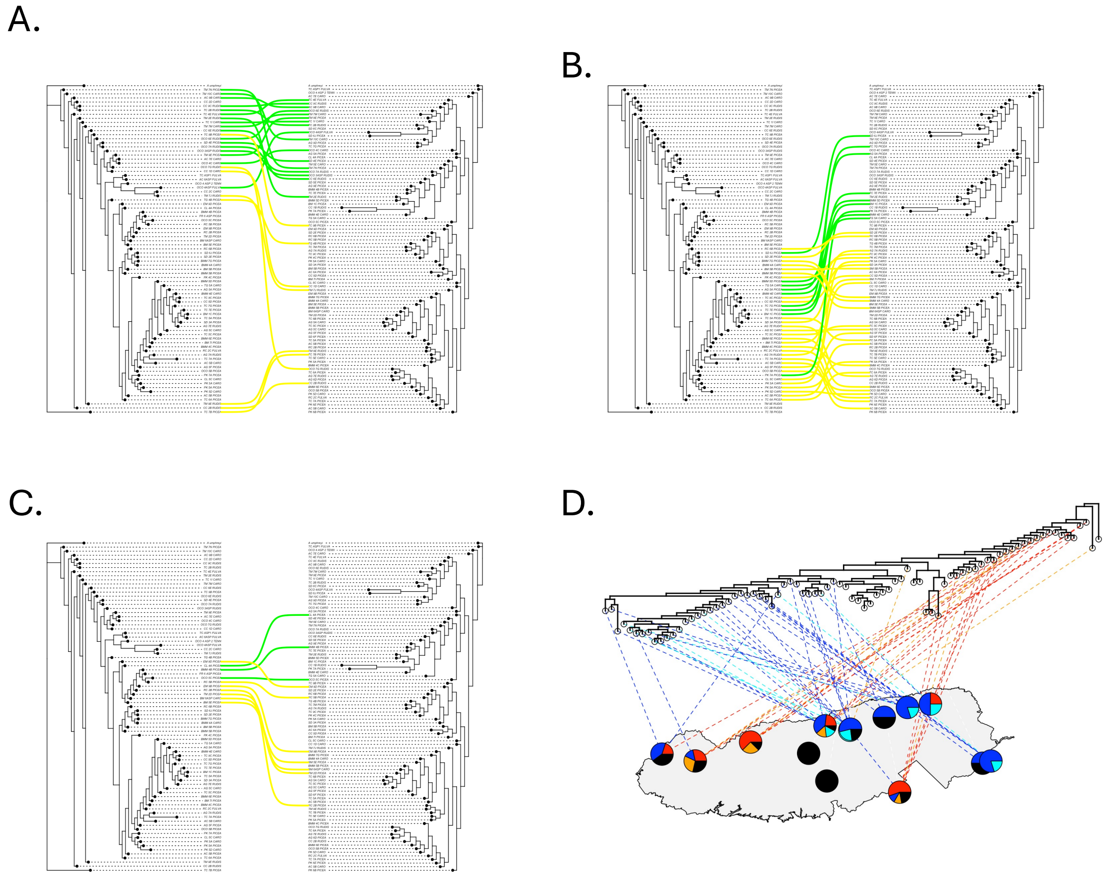


**Figure S2.1** (A-C) Tanglegrams showing how COI genes of red (A), blue (B) and purple (C) clade ants partition between the yellow and green CAD clades. (D) CO1 phylogenetic tree mapped to 14 sites of origin across 81 ants in GSMNP. The node unassociated with a GSMNP location is the outgroup, *A. umphreyi*. Lines are colored based on whether the ant is part of the red mitochondrial clade with the green nuclear clade nuclear (red), the blue mitochondrial clade with the yellow nuclear clade (blue), the red mitochondrial clade with the yellow nuclear clade (orange) the blue mitochondrial clade with the green nuclear clade (cyan) or some other combination (e.g., purple mitochondrial clade, basal nuclear 'clade').

**Table S2.1** Number of ants with blue and red COI genes and green and yellow CAD genes.

| | Green | Yellow |
|---|---|---|
| Blue | 9 | 29 |
| Red | 17 | 8 |

**Table S2.1** shows the number of ants with blue and red COI genes and green and yellow CAD genes. Blue and red clade ants differ significantly in their respective frequencies of green versus yellow CAD genes (Fisher's exact test: *p*-value = 0.0006743).

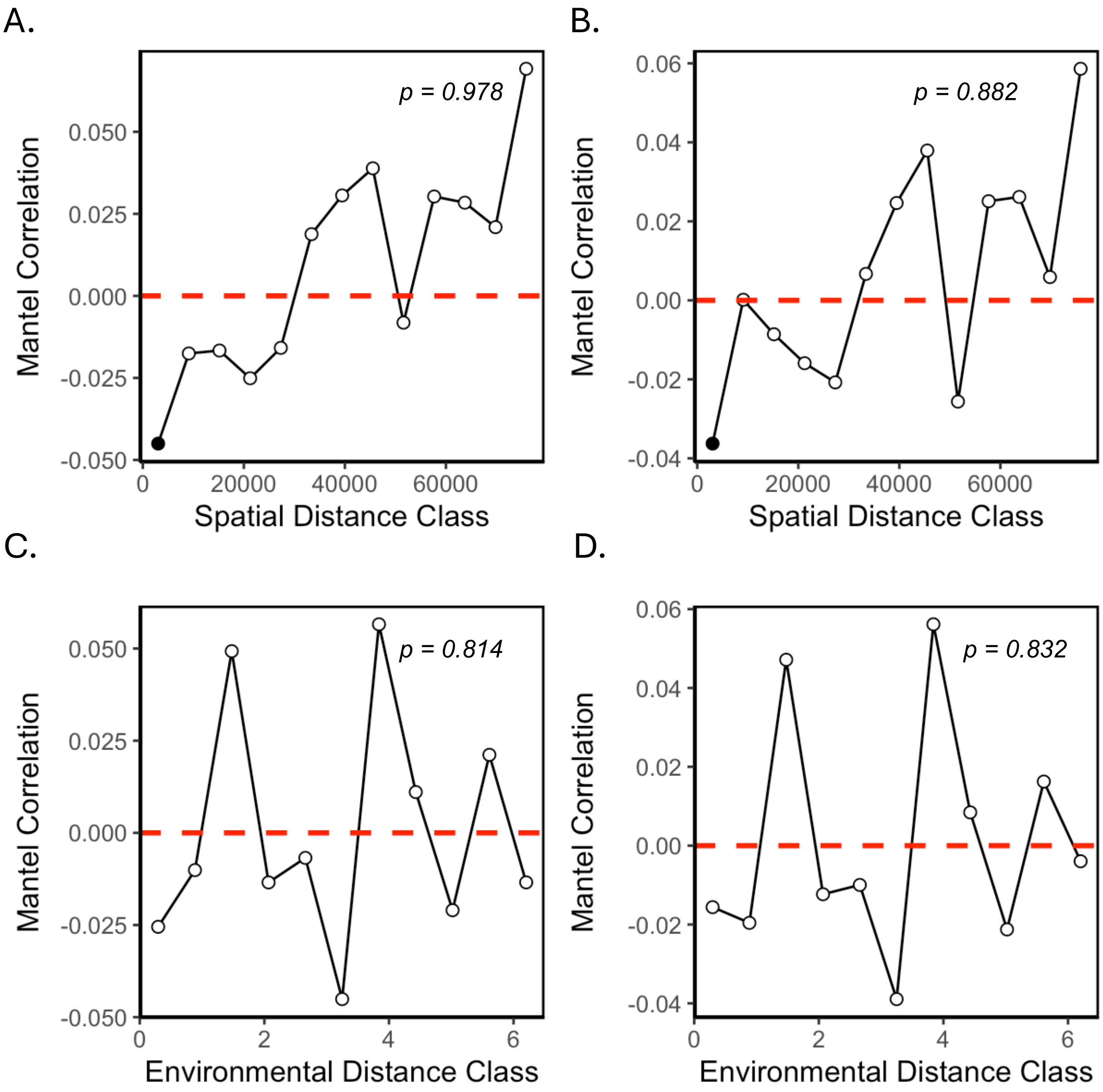


**Figure S2.2** Mantel correlograms for genetic distance based on the CAD consensus tree, regressed against spatial distance (A,B) and environmental distance (C,D) and assessed with Spearman's rank correlation (A,C) and Pearson's correlation (B,D). *p*-values shown on each panel reflect the significance of the corresponding Mantel tests.

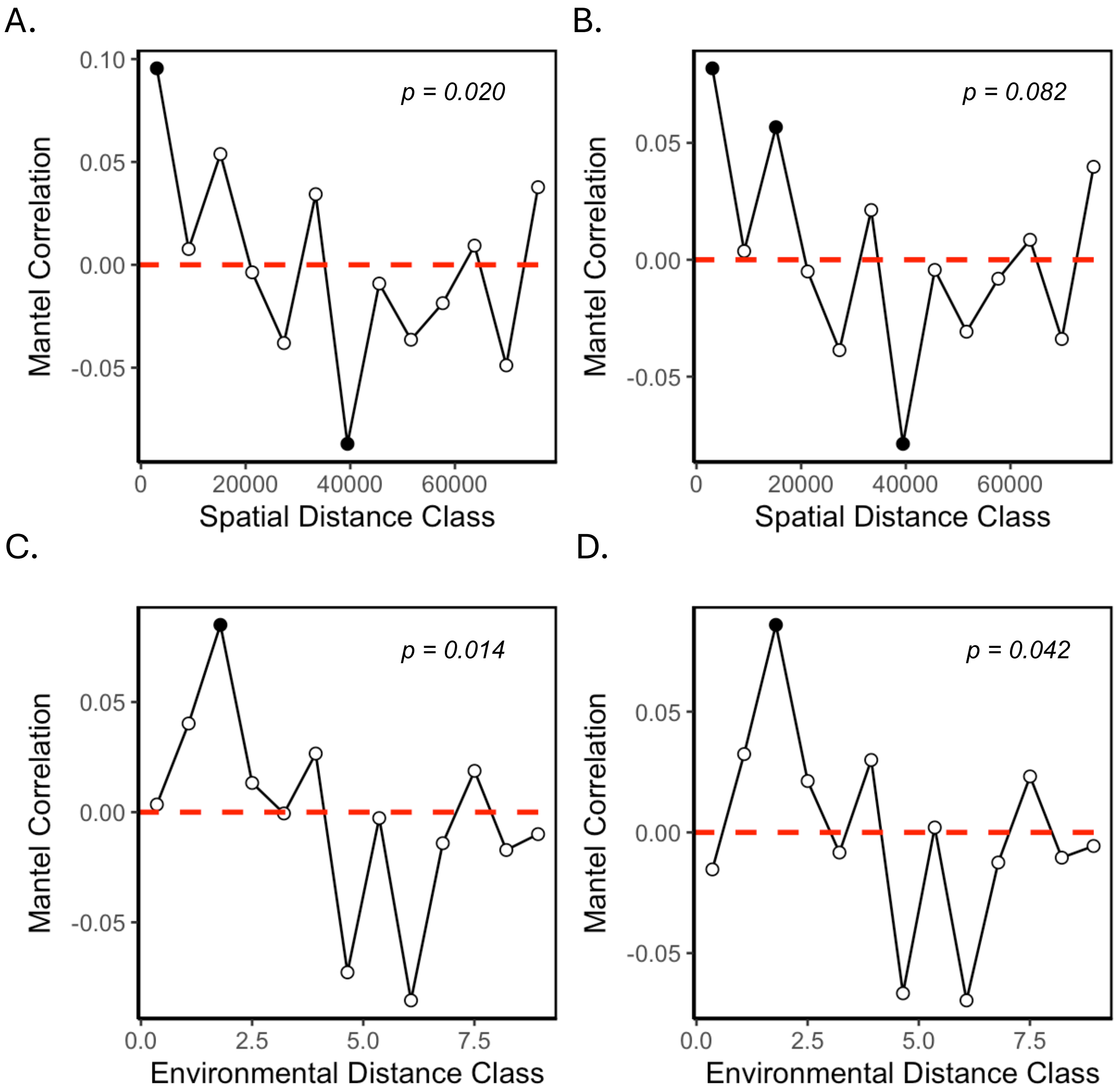


**Figure S2.3** Mantel correlograms for genetic distance based on the COI consensus tree, regressed against spatial distance (A,B) and environmental distance (C,D) and assessed with Spearman's rank correlation (A,C) and Pearson's correlation (B,D). *p*-values shown on each panel reflect the significance of the corresponding Mantel tests.

*Distance-based Redundancy Analysis*

To assess the relative contributions of spatial versus environmental factors on phylogenetic distance, we used distance-based redundancy analysis (dbRDA). Briefly, we identified distance-based Moran's eigenvector maps (MEMs) using the *dbmem* function from the *adespatial* package (version 0.3-24) to identify all MEMs corresponding to positive autocorrelation (COI: 7 MEMs, CAD: 6 MEMs). We then used these, along with our four environmental variables, as input into a dbRDA. dbRDA was performed using the *dbrda* function from the *vegan* package, regressing the phylogenetic distance matrix against MEMs and environmental variables. We then used the *vif* function from the *car* package (version 3.1-3) to sequentially remove variables with the largest variance inflation factor, until all VIF values were <3.[94] For both COI and CAD, all MEMs were retained, along with minimum soil temperature. We then ran a final dbRDA on the selected set of variables and tested for significance using the *anova.cca* function from the *vegan* package. Similar to Mantel tests, the final dbRDA for COI was highly significant (F = 3.3195, DF = 8, p-value: 0.002) whereas the final dbRDA for CAD was not (F: 1.0226, DF = 7, p-value: 0.443). Finally, we performed variance partitioning on our model, using all MEMs as our spatial variables and minimum soil temperature as our environmental variable. Variance partioning was performed using the *varpart* function from the *vegan* package. For the COI tree, spatial and spatial+environmental effects explained 20% of the variance. For the CAD tree, environmental effects explained a mere 1% of the variance, with most variance being unexplained (see Figure S2.3)

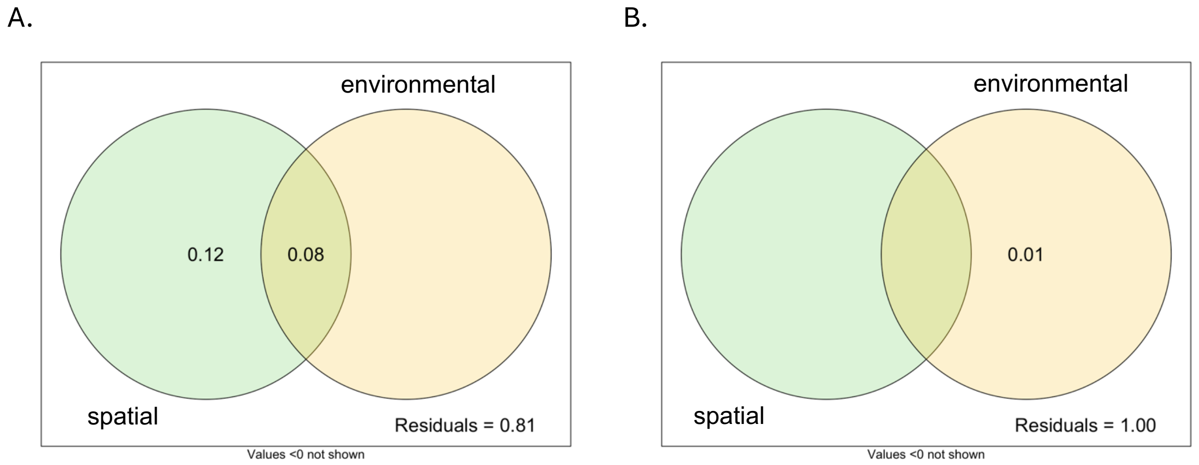


**Figure S2.4** Variance partitioning of spatial and environmental factors as explanatory variables in a dbRDA for COI phylogenetic distances (A) and CAD phylogenetic distances (B).

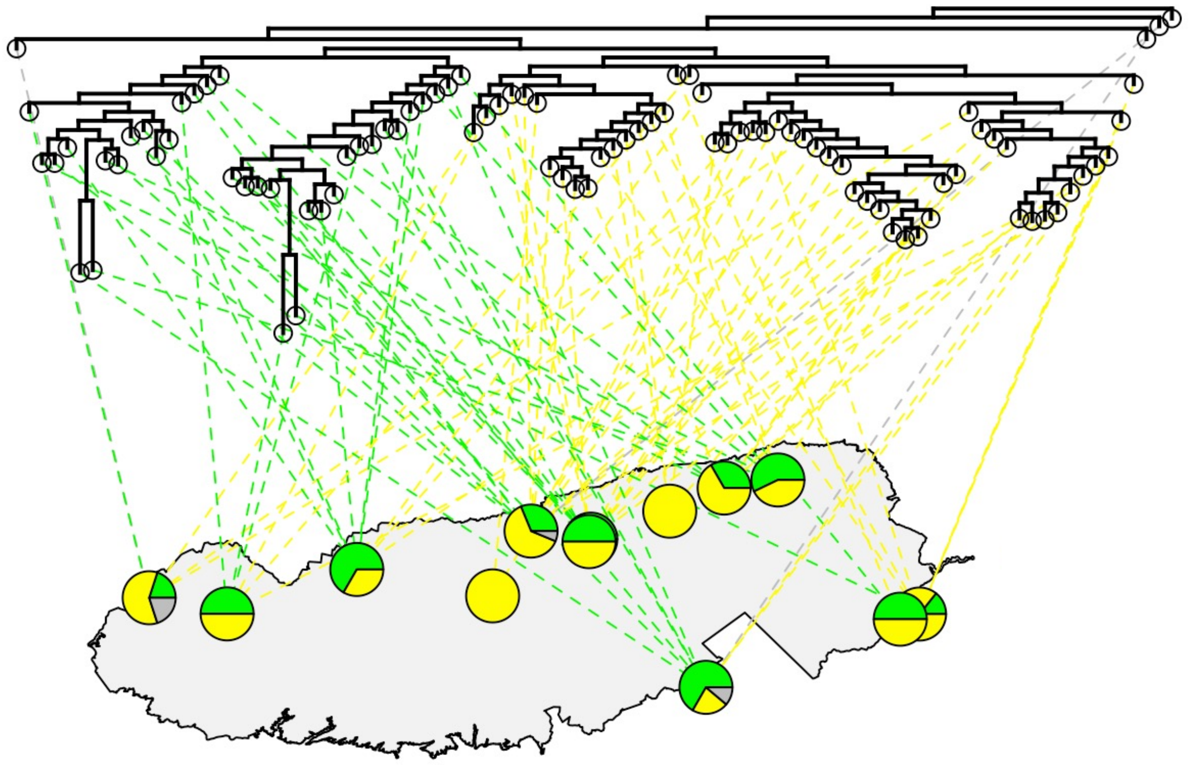

**Figure S2.5** CAD phylogenetic tree mapped to 14 sites of origin across 92 ants in GSMNP using the *plot.phylo.to.map* function in the phytools package. The node unassociated with a GSMNP location is the outgroup, *A. umphreyi*. Lines are colored based on whether the ant is included in the green or yellow clades or the basal clade (grey). Pie charts show the fraction of ants at each location that come from each clade.

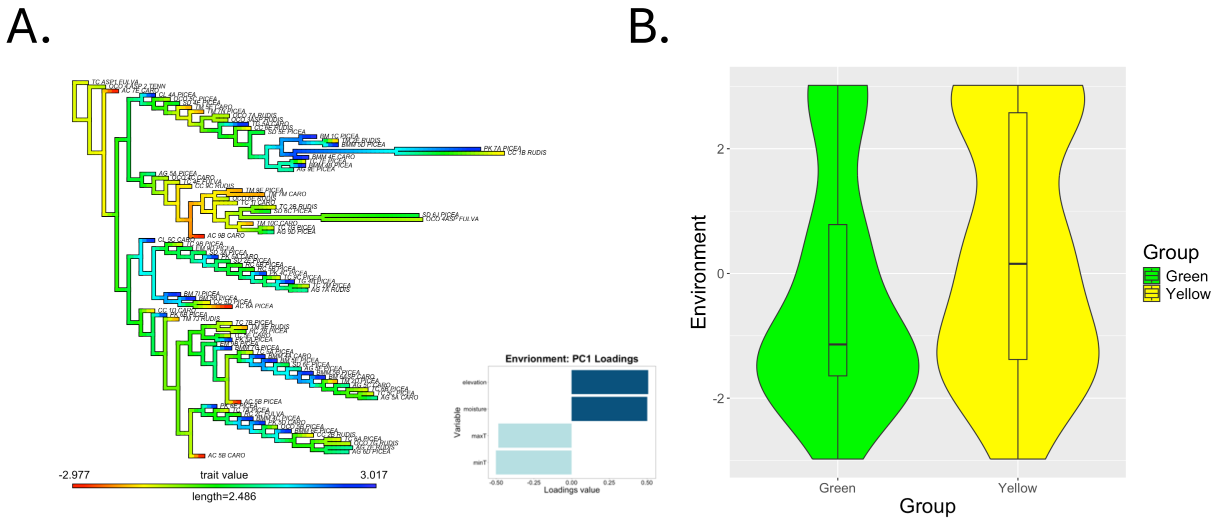


**Figure S2.6** Environmental ancestral trait reconstruction on the CAD tree based on a single ‘conglomerate’ environmental axis derived from a PCA analysis on our four environmental variables (minimum soil temperature, maximum soil temperature, soil moisture and elevation). Loadings of the four environmental variable onto the conglomerate environmental axis are shown to the left of the phylogenetic tree. **(B)** Violin plot showing comparison of values along the conglomerate environmental axis for green and yellow clade ants. Differences in environmental variables were not significant.

**Supplemental Information III: Additional Microbiota Analyses**

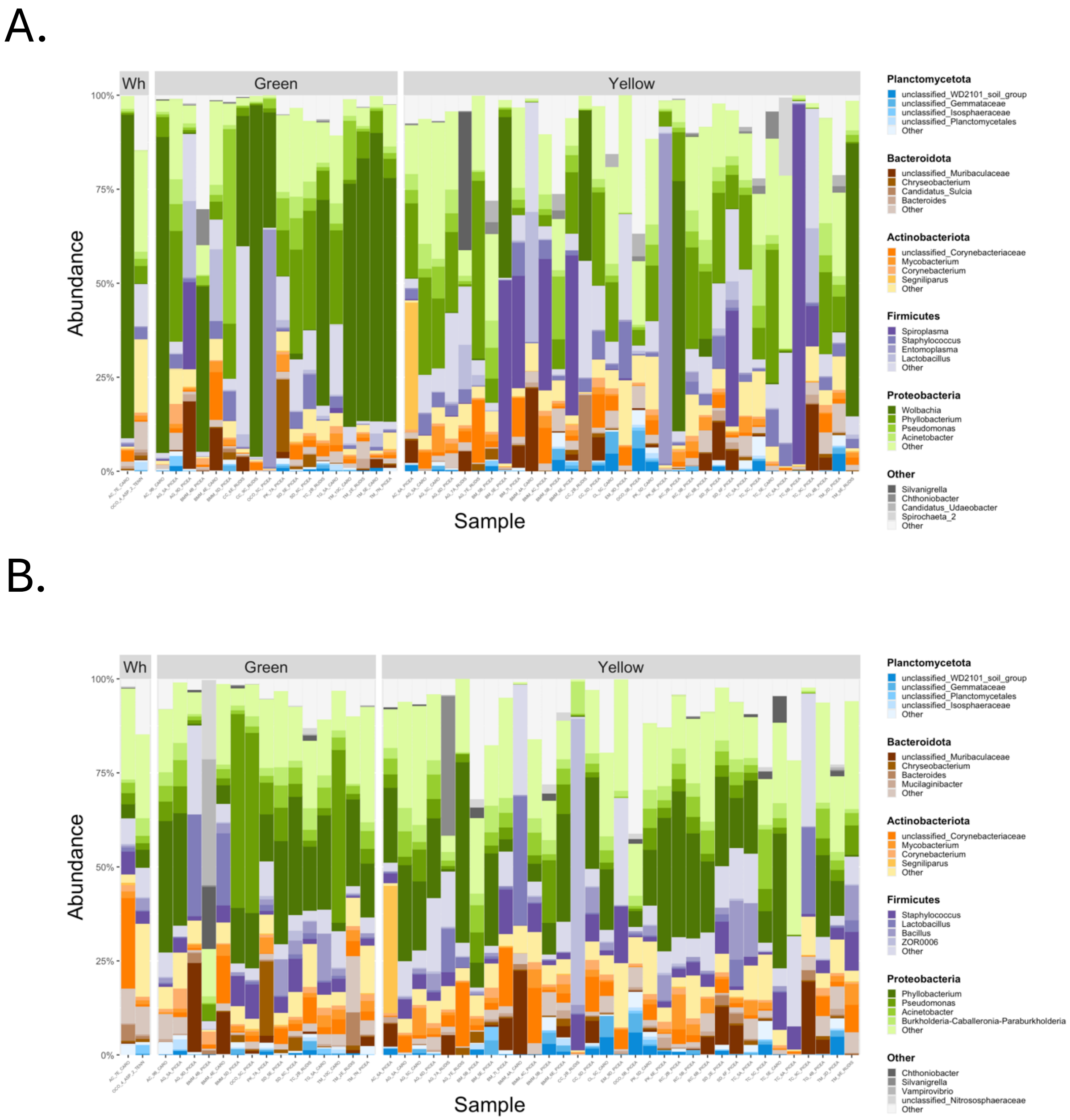


**Figure S3.1** Microbial taxon barplots for *A. rudis* complex ant microbiota with (A) and without (B) dominant endosymbiont genera (*Wolbachia*, *Spiroplasma*, *Entomoplasma*, *Sulcia*). Samples are separated according to CAD host clade and colored based on dominant phyla (Planctomycetota: blue, Bacteroidota: brown, Actinobacteriota: orange, Firmicutes: purple, Proteobacteria: green, Other: grey) and genera. Note that several samples from the red clade were dropped due to low reads following removal of endosymbionts.

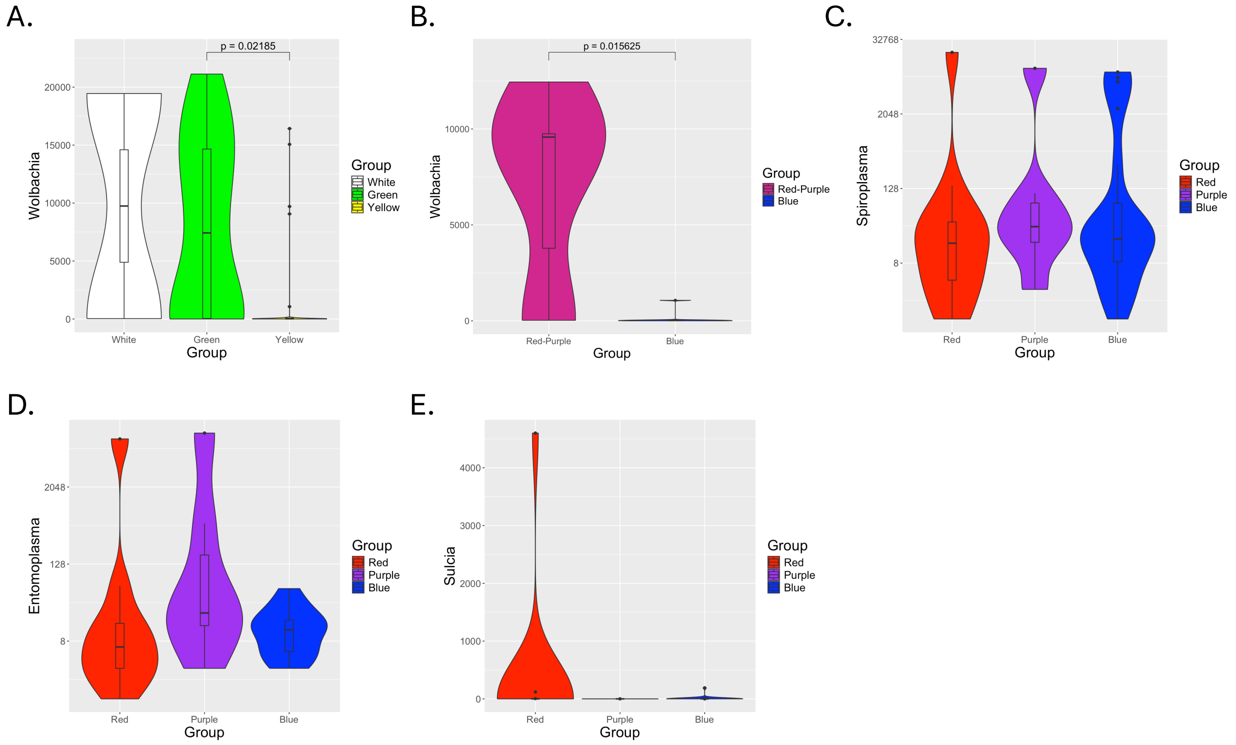


**Figure S3.2** Violin plot of (A) *Wolbachia* abundance across the two CAD clades (green and yellow respectively), along with the three basal ants (white); (B) *Wolbachia* abundance across COI clades red+purple clade ants (red-violot) and blue clade ants (blue) comparing average abundance of *Wolbachia* in each clade at each of seven sites where the clades co-occur; (C) *Spiroplasma* abundance (D) *Entomoplasma* abundance and (E) *Sulcia* abundance across the three COI clades (red, purple and blue respectively). Significant differences are indicated by brackets with associated *p*-values and are based on Kruskall-Wallis tests (A,C-E) and paired Wilcoxon sign rank tests (B).

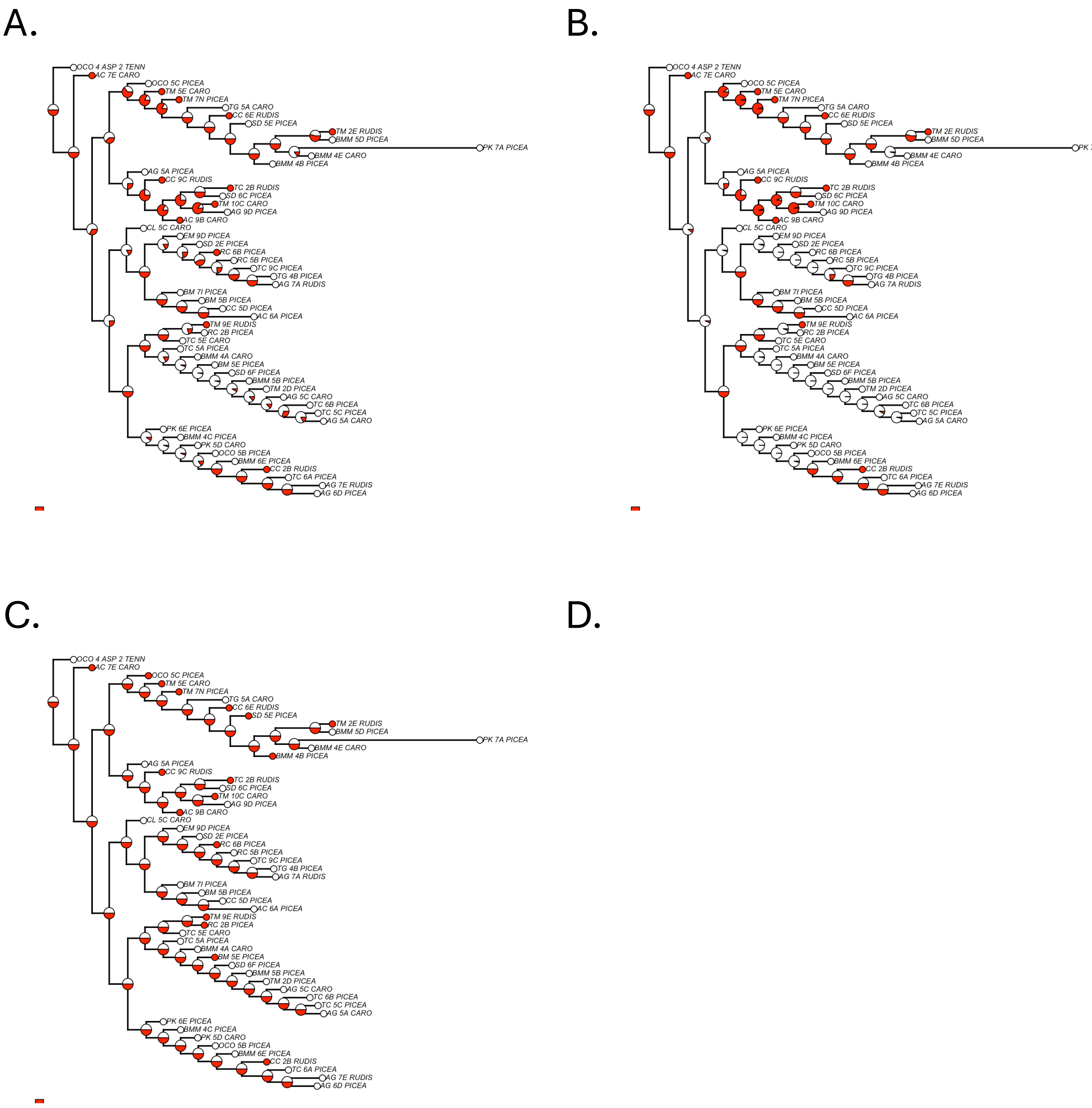


**Figure S3.3** Ancestral state reconstructions for *Wolbachia* wArudF1 **(A)**, wArudA1 **(B)** and wArudA2 **(C)** based on the CAD gene tree. **(D)** Haplotype map for CAD showing carriage of the three dominant *Wolbachia* ASVs: all three *Wolbachia* ASVs (red), wArudF1 and wArudA1 (dark pink), only wArudA2 (light pink), no *Wolbachia* (white). Black is used for animals for which we had COI sequences but where the HA microbiota did not successfully extract and/or amplify. (See SI III Figure S3.3 for *Wolbachia* ASRs and a haplotype map for the CAD gene tree).

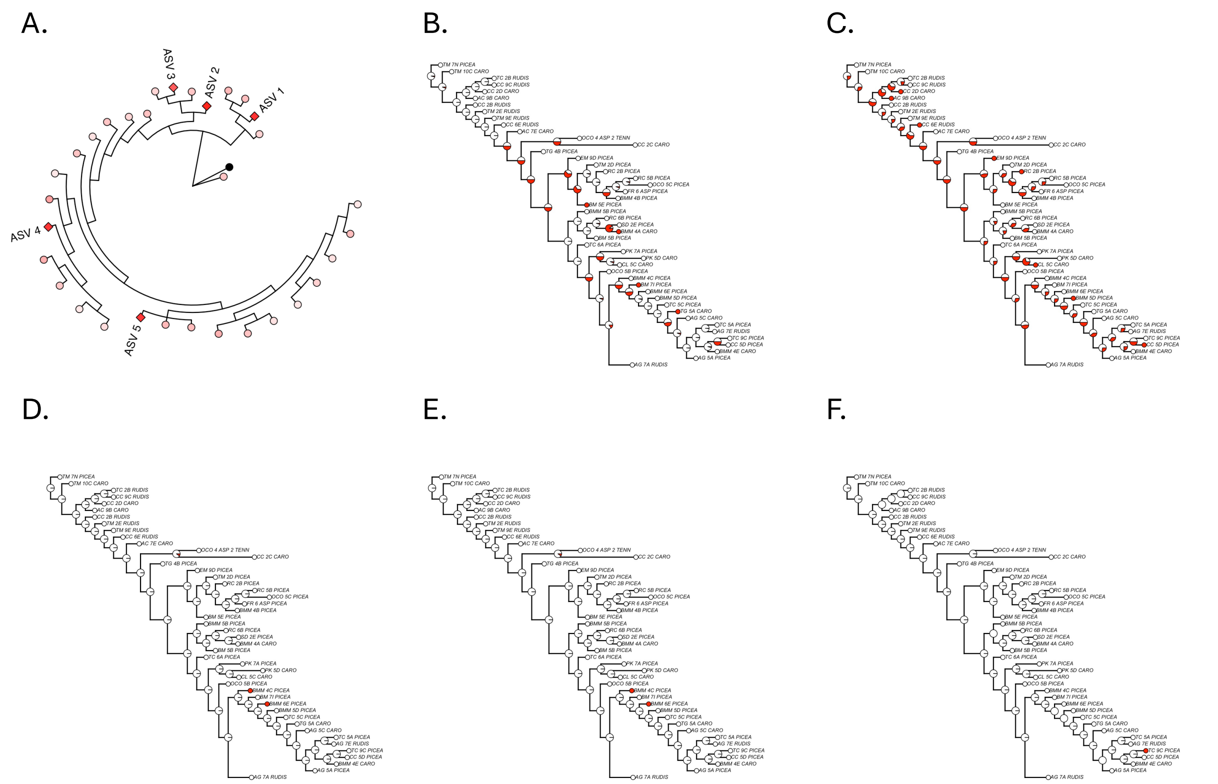


**Figure S3.4** (A) Phylogeny of the 32 *Spiroplasma* ASVs in our study, rooted with an *Entomoplasma* ASV as the outgroup (black). *Spiroplasma* tips are colored according to their overall abundance across all ant samples (red: high, white: low). Circular tips are used for non-focal ASVs and diamond tips are used for focal ASVs. Note that the color scheme is logarithmic. (B-F) Ancestral state reconstructions for *Spiroplasma* ASV1 (B), ASV2 (C), ASV3 (D), ASV4 (E) and ASV5 (F) based on the COI gene tree.

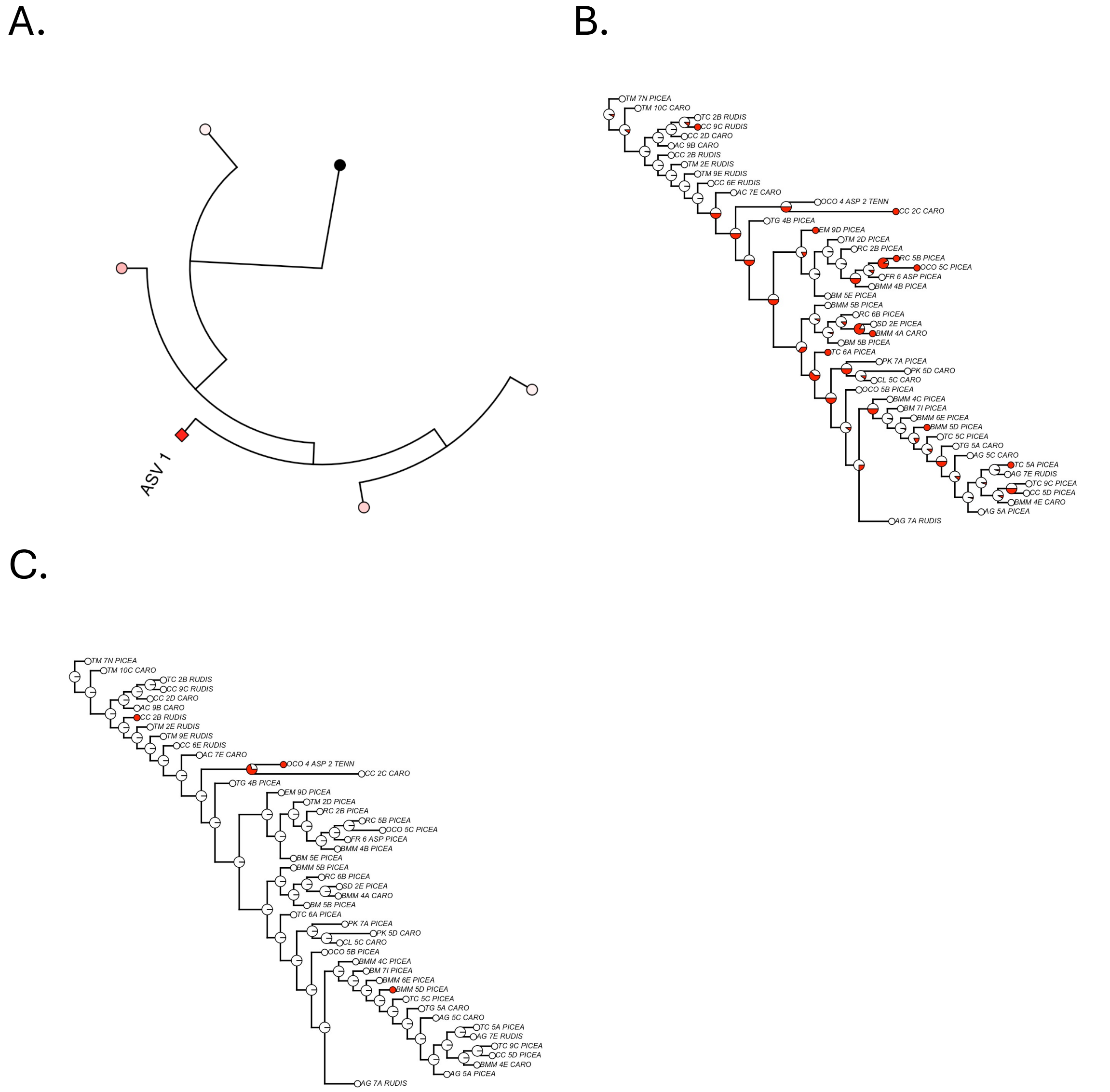


**Figure S3.5** (A) Phylogeny of the 5 *Entomoplasma* ASVs in our study, rooted with an *Spiroplasma* ASV as the outgroup (black). *Entomoplasma* tips are colored according to their overall abundance across all ant samples (red: high, white: low). Circular tips are used for non-focal ASVs and diamond tips are used for focal ASVs. Note that the color scheme is logarithmic. (B-F) Ancestral state reconstructions for *Entomoplasma* ASV1 (B) and *Sulcia* ASV1 (C).

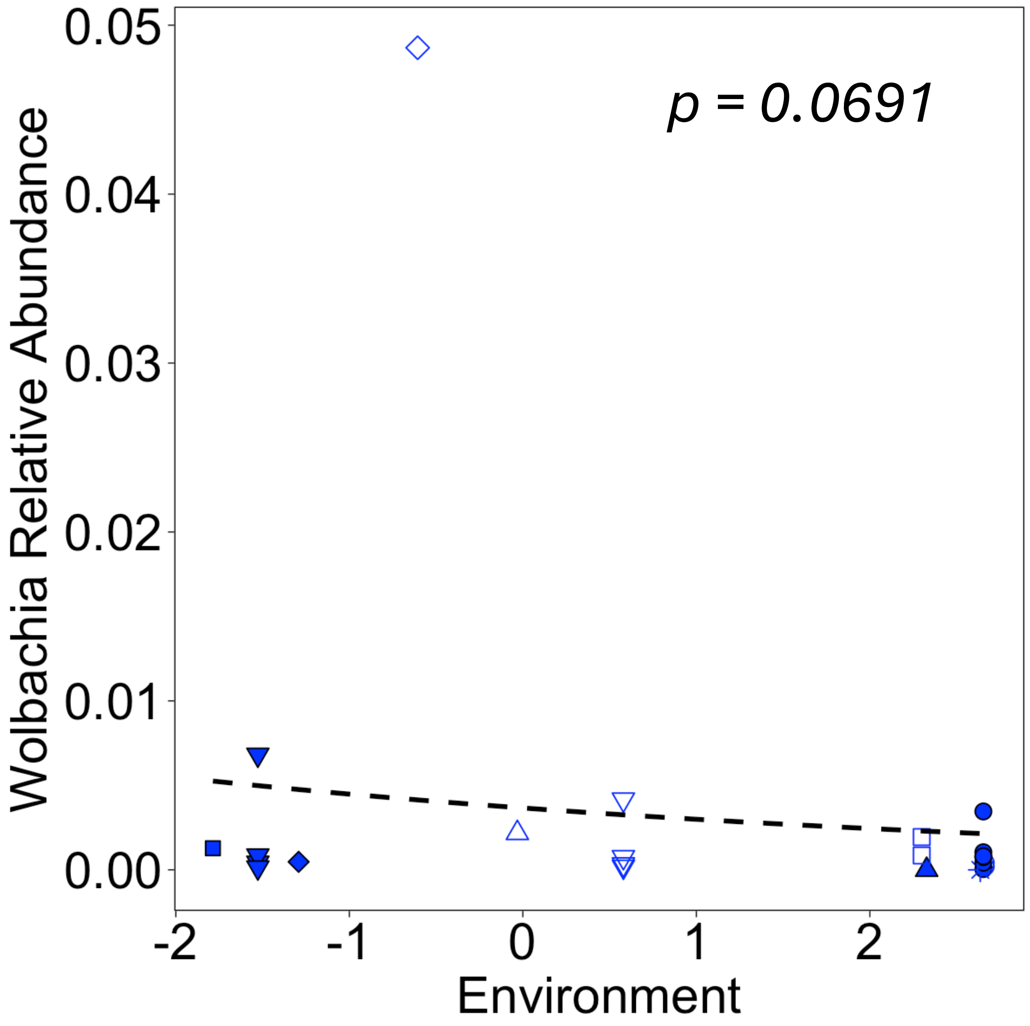


**Figure S3.6** Generalized linear models of *Wolbachia* relative abundance as a function of our conglomerate environmental variable (see Methods) for blue clade ants including the outlier at Ramsey Cascade. Sites are denoted as follows: Cades Cove (closed squares), Twin Creeks (closed downward triangles), Oconaluftee (closed diamond), Trillium Gap (star), Snake Den (open upward triangle), Albright Grove (open downward triangle), Cataloochee (closed upward blue triangles), Purchase Knob (open squares), Brush Mountain Myrtle (closed blue circles), Brush Mountain (open circles), Ramsey Casdae (open diamond).

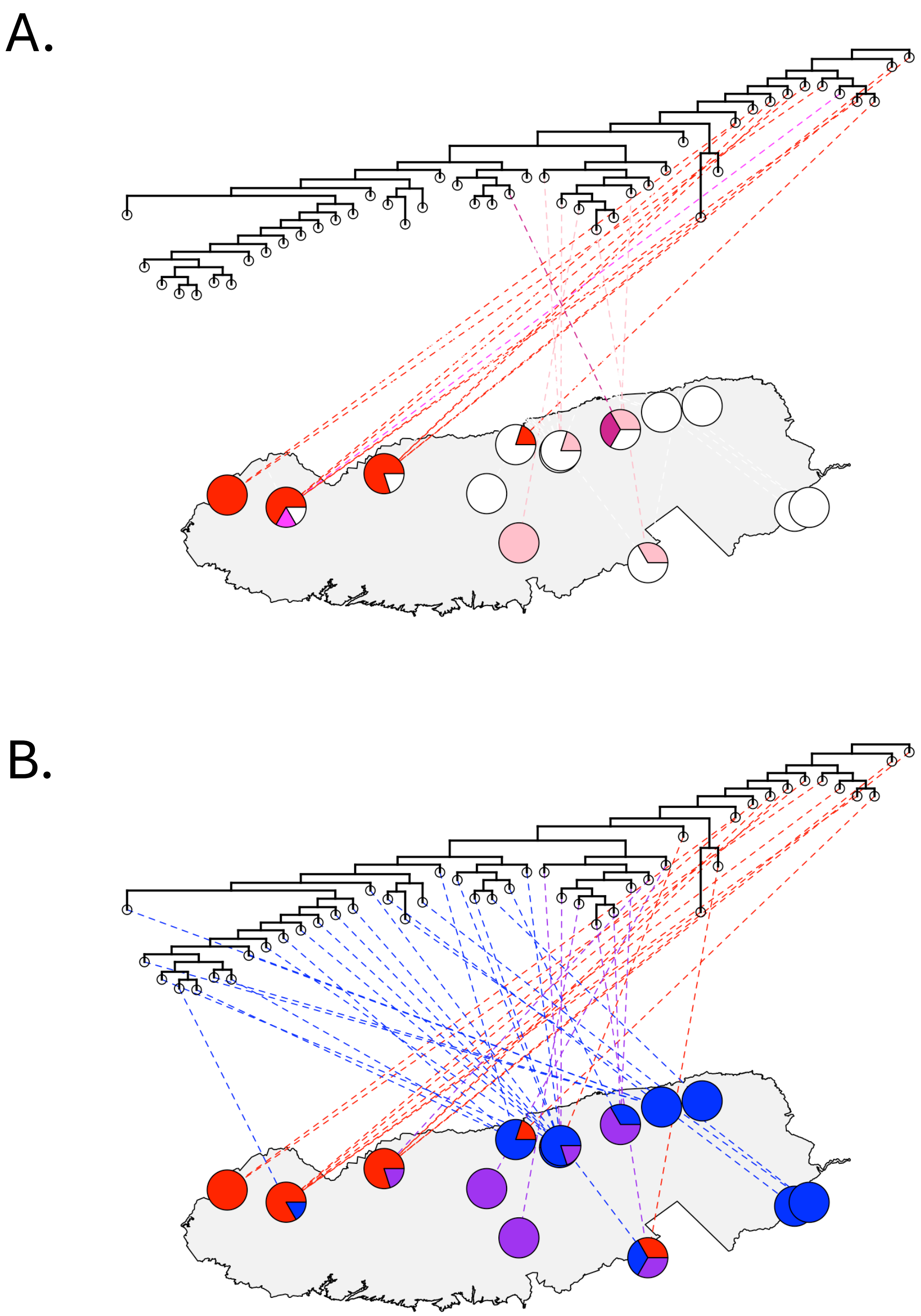


**Figure S3.7** CO1 phylogenetic tree mapped to 14 sites of origin across 81 ants in GSMNP. (A) Lines are colored based on *Wolbachia* carriage as follows: wArudF1+wArudA1+wArudA2 (red), wArudA1+wArudA2 (bright pink), wArudF1+wArudA1 (dark pink), wArudA2 (pink), and no *Wolbachia* (white). (B) Lines are colored based on COI clade. Note that in (B), to facilitate comparison with the *Wolbachia* map, we only include COI sequences for which we have paired HA microbiota sequences.

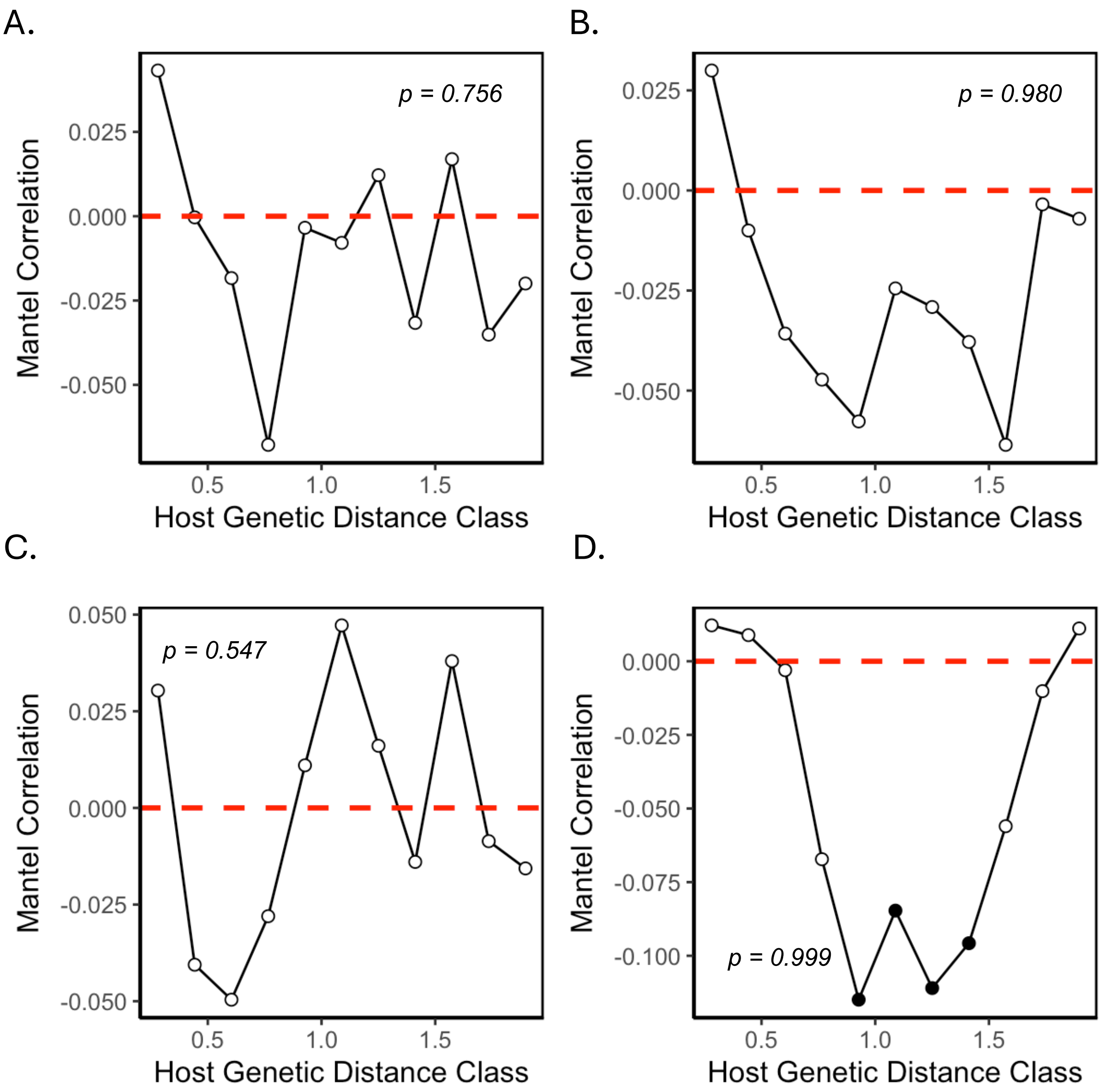


**Figure S3.8** Mantel correlograms for HA microbiota (with endosymbionts removed) Jaccard (A), Bray-Curtis (B), UniFrac (C) and weighted UniFrac (D) distances regressed against host genetic distance based on the COI and assessed with Spearman's correlation. *p*-values shown on each panel reflect the significance of the corresponding Mantel tests.

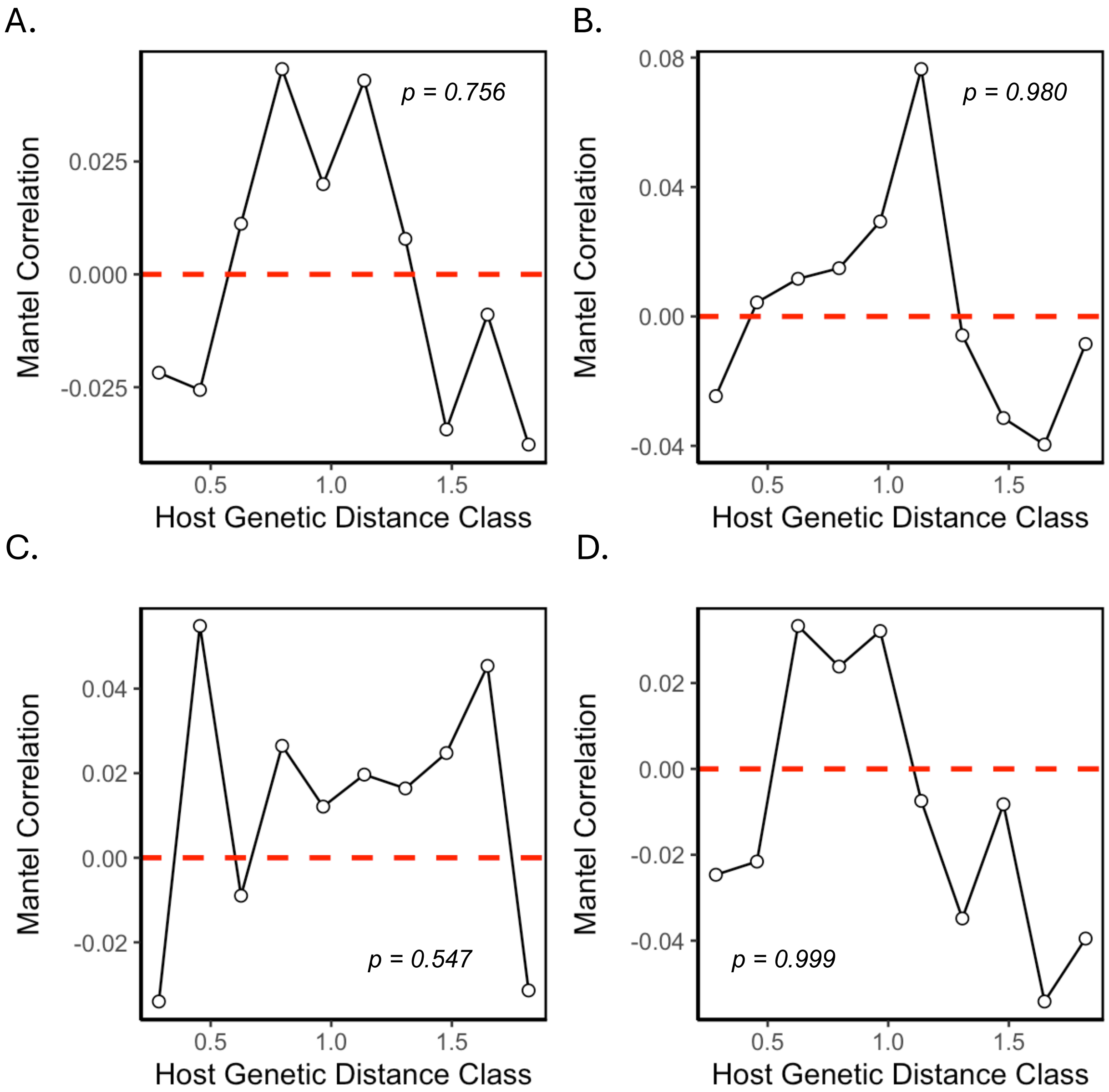


**Figure S3.9** Mantel correlograms for HA microbiota (with endosymbionts removed) Jaccard (A), Bray-Curtis (B), UniFrac (C) and weighted UniFrac (D) distances regressed against host genetic distance based on the CAD and assessed with Spearman's correlation. *p*-values shown on each panel reflect the significance of the corresponding Mantel tests.

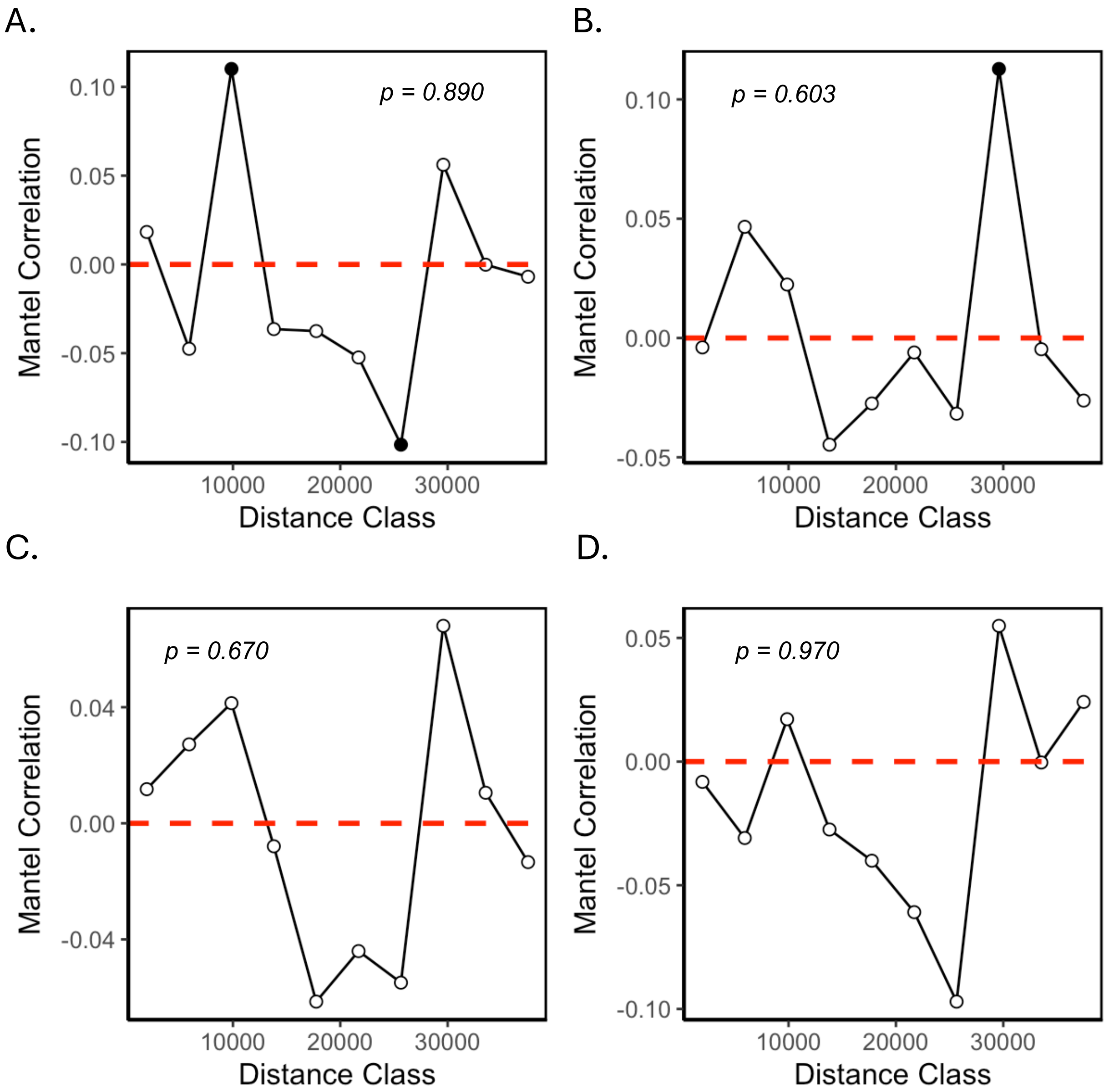


**Figure S3.10** Mantel correlograms for HA microbiota similarity (with endosymbionts removed) using Jaccard (A), Bray-Curtis (B), UniFrac (C) and weighted UniFrac (D) distances regressed against spatial distance and assessed with Spearman's correlation. *p*-values shown on each panel reflect the significance of the corresponding Mantel tests.

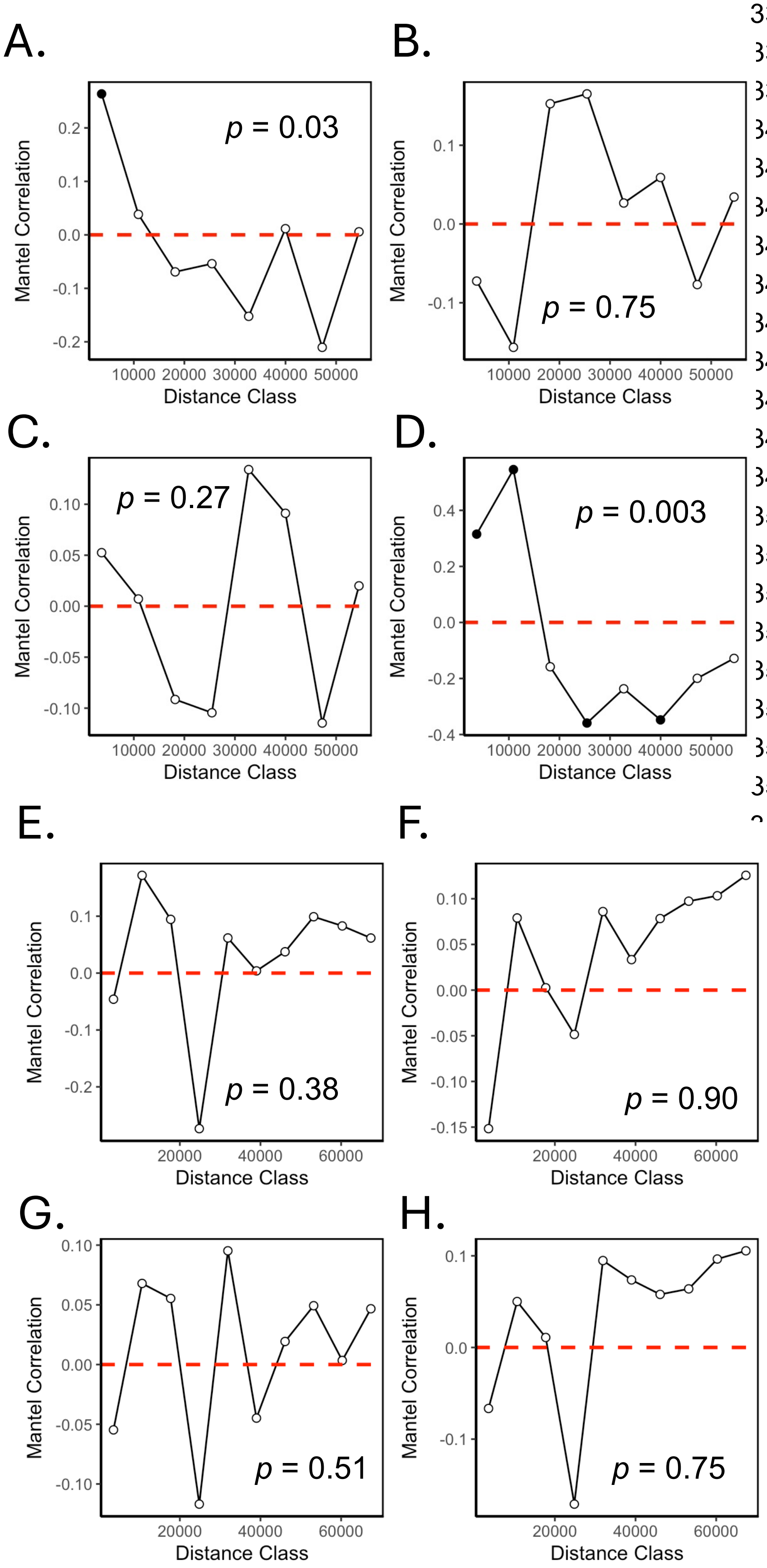


**Figure S3.11** Mantel correlograms for HA microbiota (with endosymbionts removed) similarity of red (A-D) and blue (E-H) clade ants using Jaccard (A,E), Bray-Curtis (B,F), UniFrac (C,G) and weighted UniFrac (D,H) distances regressed against spatial distance and assessed with Spearman's correlation. *p*-values shown on each panel reflect the significance of the corresponding Mantel tests.

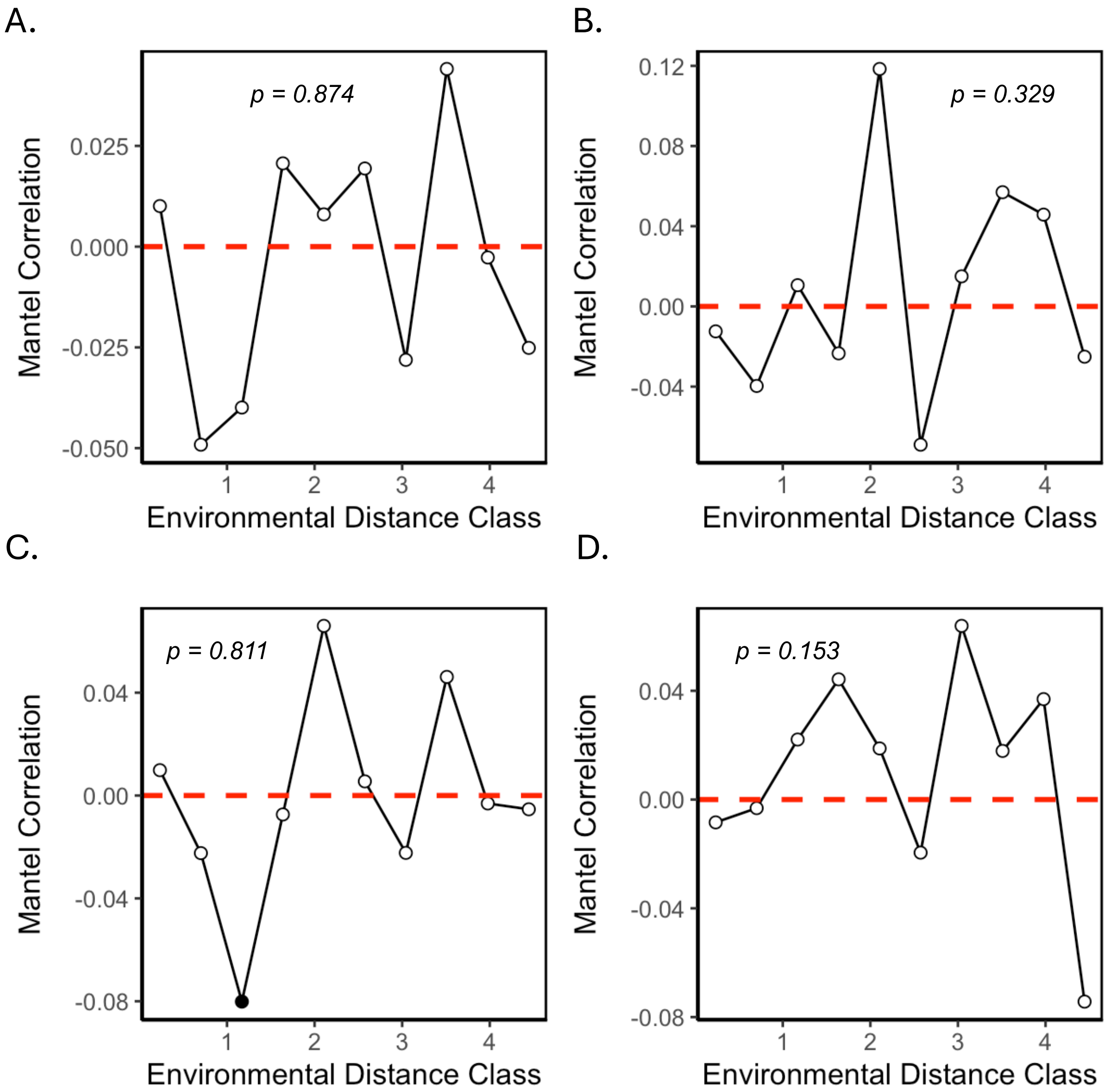


**Figure S3.12** Mantel correlograms for HA microbiota (with endosymbionts removed) Jaccard (A), Bray-Curtis (B), UniFrac (C) and weighted UniFrac (D) distances regressed against environmental distance using Spearman's correlation. *p*-values shown on each panel reflect the significance of the corresponding Mantel tests.

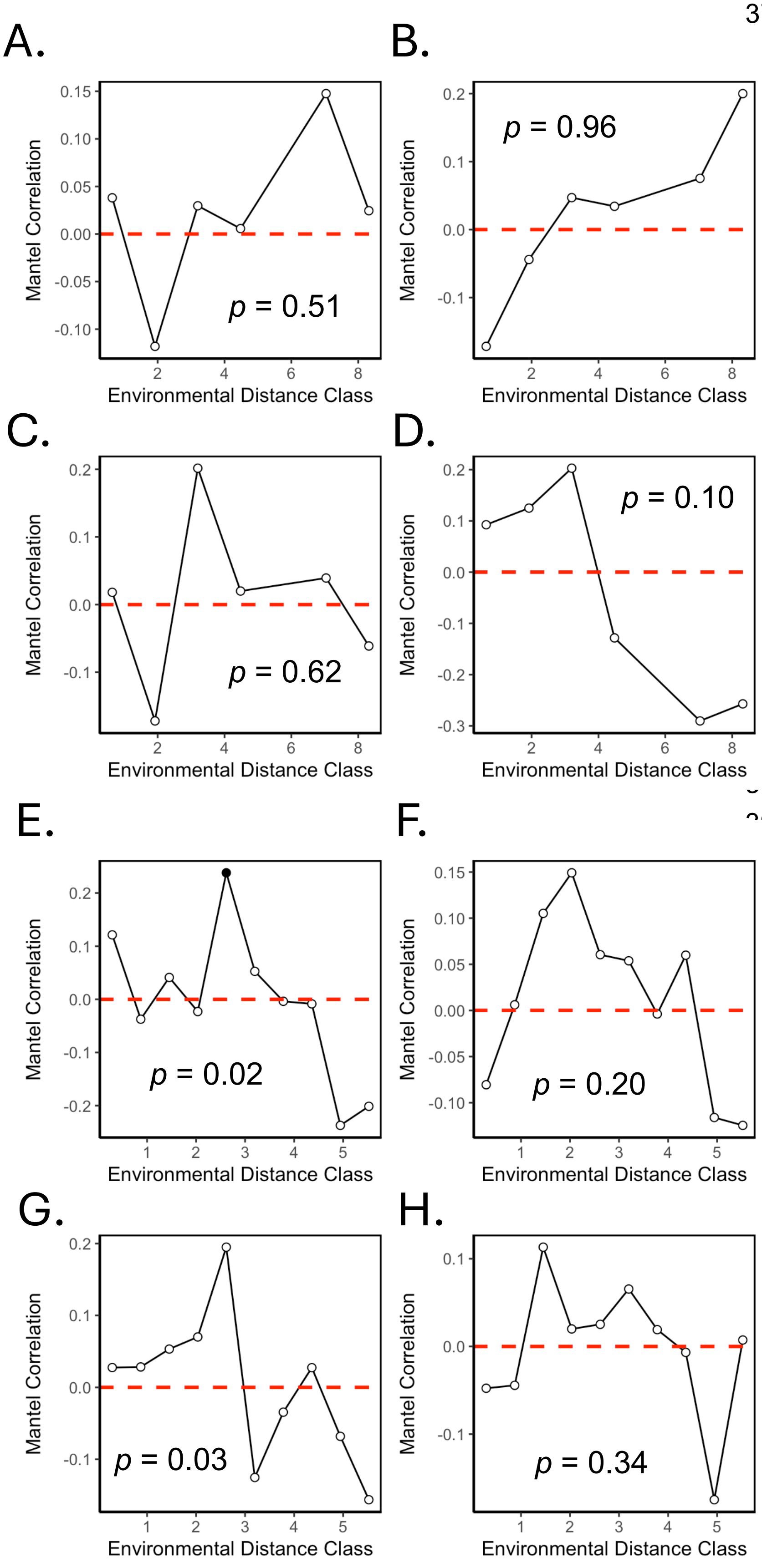


**Figure S3.13** Mantel correlograms for HA microbiota (with endosymbionts removed) similarity of red (A-D) and blue (E-H) clade ants using Jaccard (A,E), Bray-Curtis (B,F), UniFrac (C,G) and weighted UniFrac (D,H) distances regressed against environmental distance and assessed with Spearman's correlation. *p*-values shown on each panel reflect the significance of the corresponding Mantel tests.

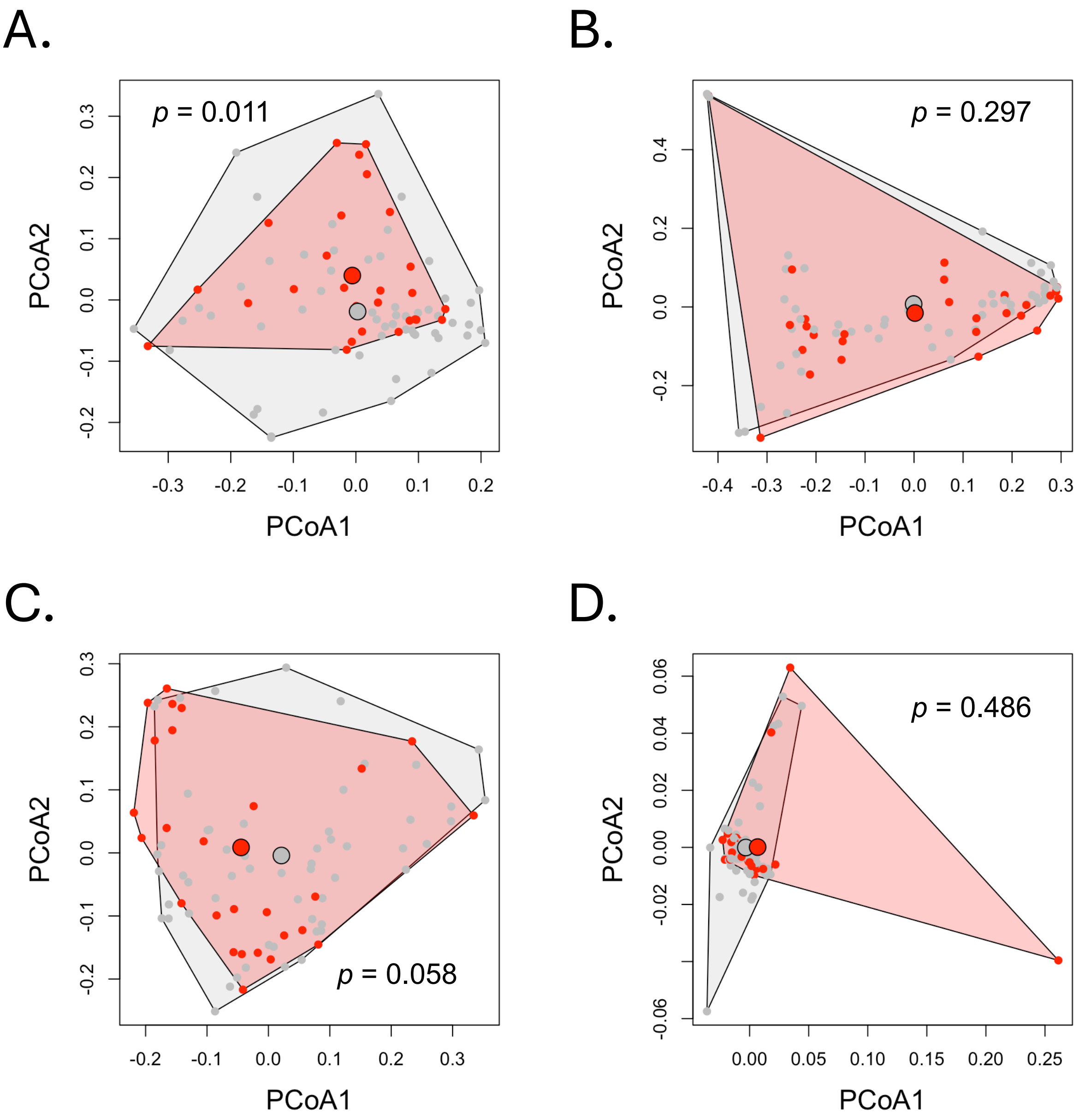


**Figure S3.14** Principal Coordinates Analyses based on Jaccard (A), Bray-Curtis (B), UniFrac (C) and weighted UniFrac (D) distances showing microbiota (endosymbionts removed) from ants that, prior to removal of endosymbionts, had microbiota comprising >0.5% (red) and <0.5% (grey) *Wolbachia*. *p*-values shown on each panel reflect the significance of the corresponding PERMANOVA tests.

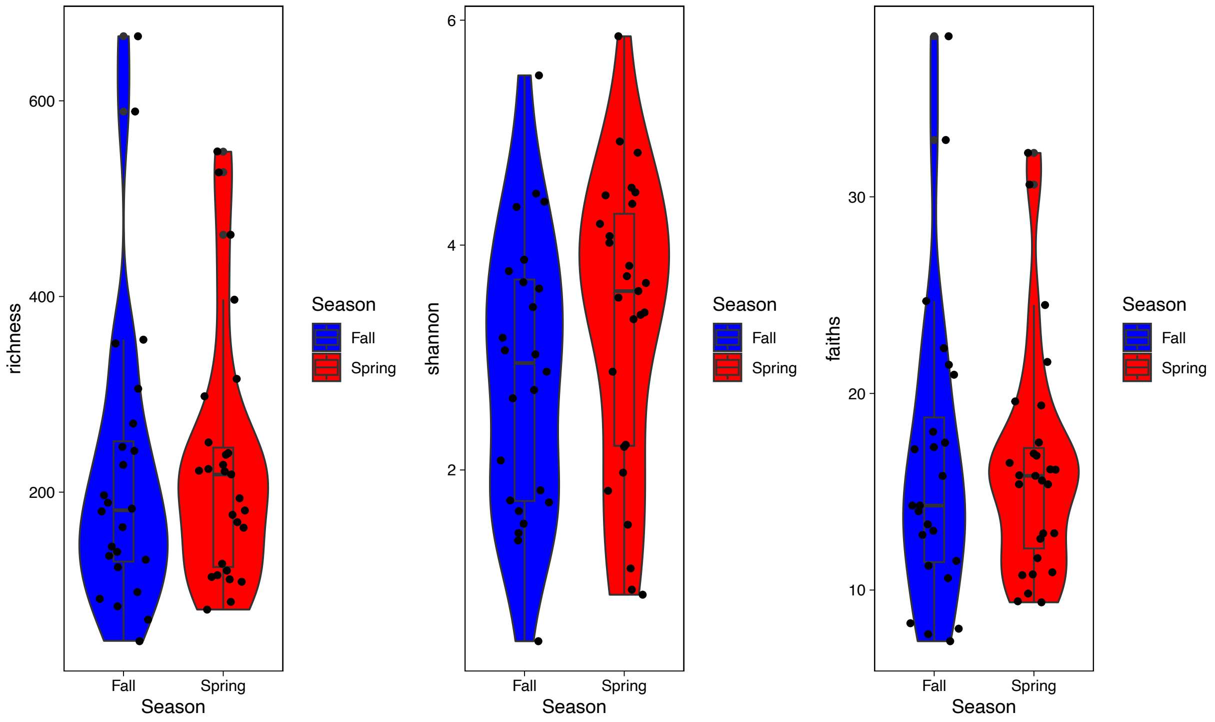


**Figure S3.15** Comparison of alpha diversity between Spring and Summer samples. Richness did not differ by season (W = 301, p-value = 0.6711). Shannon diversity did not differ by season (W = 248, p-value = 0.1554). Faith's PD did not differ by season (W = 316, p-value = 0.8884).

**Table S3.1** Summary of results for Partial Mantel tests and dbRDA on Microbiota similarity regressed against spatial distance and environment.

| | Partial Mantel | | dbRDA | |
|---|---|---|---|---|
| | statistic | p-value | statistic | p-value |
| | Jaccard | | | |
| All | -0.02739 | 0.678 | 1.0312 | 0.181 |
| Red Clade | -0.1406 | 0.762 | 1.1672 | 0.014* |
| Blue Clade | 0.2448 | 0.009** | 0.9594 | 0.777 |
| | Bray-Curtis | | | |
| All | 0.03281 | 0.281 | 1.0292 | 0.354 |
| Red Clade | -0.2349 | 0.945 | 0.9337 | 0.574 |
| Blue Clade | 0.1275 | 0.079 | 0.9013 | 0.723 |
| | UniFrac | | | |
| All | -0.03193 | 0.749 | 1.0041 | 0.425 |
| Red Clade | -0.0764 | 0.633 | 0.9828 | 0.576 |
| Blue Clade | 0.1631 | 0.012* | 0.9996 | 0.457 |
| | weighted UniFrac | | | |
| All | 0.1374 | 0.028* | 0.9463 | 0.428 |
| Red Clade | 0.2874 | 0.215 | 1.9006 | 0.016* |

| | | | | |
|---|---|---|---|---|
| Blue Clade | 0.05287 | 0.275 | 0.6462 | 0.897 |
